# Partial suppression of BCAA catabolism as a potential therapy for BCKDK deficiency

**DOI:** 10.1101/2023.10.12.560929

**Authors:** Laura Ohl, Amanda Kuhs, Ryan Pluck, Emily Durham, Michael Noji, Nathan D Philip, Zoltan Arany, Rebecca C Ahrens-Nicklas

## Abstract

Branched chain ketoacid dehydrogenase kinase (BCKDK) deficiency is a recently described inherited neurometabolic disorder of branched chain amino acid (BCAA) metabolism implying increased BCAA catabolism. It has been hypothesized that a severe reduction in systemic BCAA levels underlies the disease pathophysiology, and that BCAA supplementation may ameliorate disease phenotypes. To test this hypothesis, we characterized a recent mouse model of BCKDK deficiency and evaluated the efficacy of enteral BCAA supplementation in this model. Surprisingly, BCAA supplementation exacerbated neurodevelopmental deficits and did not correct biochemical abnormalities despite increasing systemic BCAA levels. These data suggest that aberrant flux through the BCAA catabolic pathway, not just BCAA insufficiency, may contribute to disease pathology. In support of this conclusion, genetic re-regulation of BCAA catabolism, through *Dbt* haploinsufficiency, partially rescued biochemical and behavioral phenotypes in BCKDK deficient mice. Collectively, these data raise into question assumptions widely made about the pathophysiology of BCKDK insufficiency and suggest a novel approach to develop potential therapies for this disease.

## 1. Introduction

Branched-chain amino acids (BCAA), leucine, isoleucine, and valine, are essential amino acids, required for cellular function and growth. Abnormalities of BCAA metabolism have been implicated in a variety of human diseases including inborn errors of metabolism, diabetes, cancer, heart failure, and neurodegenerative diseases^1,2^. BCAA levels are controlled by a highly regulated catabolic pathway. In the first step of the pathway, BCAAs are converted into alpha-keto acids by a branched-chain aminotransferase. These alpha-keto acids are then further broken down by the rate limiting enzyme branched-chain ketoacid dehydrogenase complex (BCKDH). The BCKDH complex and subsequent downstream enzymes convert the alpha-keto acids into intermediates, including acetyl-CoA and succinyl-CoA. These CoA species can feed into the tricyclic acid (TCA) cycle, which is coupled to the electron transport chain in the mitochondria. Within the mitochondria, branched-chain keto-acid dehydrogenase kinase (BCKDK) phosphorylates BCKDH and inhibits its activity to prevent excessive catabolism of BCAAs and BCKAs^2^.

BCAAs cross the blood brain barrier and regulate several key processes in the central nervous system. BCAAs provide approximately 30% of the nitrogen groups required for the synthesis of glutamate, the major excitatory neurotransmitter of the brain^3^. As such, BCAAs provide essential precursors for neurotransmitter biosynthesis and recycling. Additionally, BCAAs serve as key energy sources, as they donate their carbon skeleton to TCA cycle intermediates when catabolized^3^. Although the biochemistry of BCAA catabolism is well studied, how perturbation of this pathway leads to brain dysfunction is not well understood.

Pathogenic variants in BCKDK lead to a syndrome characterized by intellectual disability, postnatal microcephaly, autism, and seizures in human patients^4^. To date, there are 14 reported unique pathogenic variants located throughout the protein structure^4–6^. BCKDK deficiency patients all have reduced BCAA levels in the blood, which has resulted in BCAA supplementation being proposed as a potential therapy^4^. However, recent studies of high protein diets with BCAA supplementation and its impact on brain function reveal nominal improvement of neurodevelopmental skills and neurodevelopment^6,7^.

Furthermore, how BCKDK pathogenic variants induce neurological symptoms remains understudied. Previous studies of BCKDK deficiency have focused on behavioral characterization of mouse models, amino acid level changes in mice and humans, and mitochondrial defects observed in patient-derived fibroblasts^4,8,9^. However, an explanation as to how these changes alter human brain function remains to be elucidated.

Here, we investigate the consequences of loss of BCKDK function in the brain by utilizing a recently published *Bckdk*^-/-^ mouse model^10^. In addition, we evaluate if implementation of BCAA repletion therapy in the postnatal period can prevent early neurodevelopmental deficits and postnatal microcephaly. Finally, we compare BCAA supplementation to an alternative therapeutic approach, re-regulating BCAA catabolism through genetic inhibition.

## 2. Materials and Methods

### 2.1. Animals

All experiments with animals were approved by the Institutional Animal Care and Use Committee at the Children’s Hospital of Philadelphia and guidelines set by the US Public Health Service’s Policy on Humane Care and Use of Laboratory Animals. *Bckdk*^-/-^ Mice were generated, published, and gifted from the Arany lab.^10^ Mice were collected after cryoanesthetization (p0) followed by decapitation or euthanasia with isoflurane (p21) and cervical dislocation followed by decapitation. Heads were collected for µCT analysis for microcephaly at p0 and p21. Brains and livers were collected, and flash frozen for metabolic analysis at p21. Mice were perfused with PBS followed by 4% PFA for representative images of brains.

*Dbt ^-/+^* ^mice^ were generated from a novel floxed *DBT* allele mouse on the C57BL/6J background. *Dbt ^Flox/Flox^* mice with loxP sites flanking exon 2 of *Dbt* were generated by the University of Pennsylvania CRISPR/Cas9 Mouse Targeting Core using the Alt-R™ CRISPR-Cas 9 genome editing system (Integrated DNA Technologies, Coralville, IA). The target sequences were as follows: 5′ - GTGAAGGTTATTGACAGAGTGGG; 3′ - CATTGTGGGAAACTATGGAGCGG. CRISPR guides were designed, and the crispr RNA (crRNA) sequences were as follows: 5′ - GTGAAGGTTATTGACAGAGT; 3′ - CATTGTGGGAAACTATGGAG. The trans-activating crRNA (tracrRNA, Integrated DNA Technologies, #1072533) then hybridizes to crRNA to activate the Cas9 enzyme. After oocyte injection, founders were confirmed via Sanger sequencing of genomic DNA. Founders were backcrossed to C57BL/6J mice, which were then bred to homozygosity (*Dbt Flox/Flox*).

*Dbt ^Flox/Flox^* mice were then crossed with *Nestin ^Cre+^* mice (The Jackson Laboratory, Bar Harbor, ME; B6.Cg-Tg(Nes-cre)1Kln/J, Stock No: 003771), which express Cre in both the nervous system and oocytes, resulting in the frequent germline recombination of Cre in pups^31^. Offspring that demonstrated germline recombination of the floxed allele were then bred to *Dbt^+/+^* mice to produce *Dbt ^+/-^* (*Nestin ^Cre-^*) mice, which were used for breeding.

### 2.2. Genotyping

Mouse tail snips were obtained for genotyping. DNA extraction was performed using the DNeasy Blood & Tissue Kit (QIAGEN, Hilden, Germany, #69506).

For *Bckdk* genotyping, a combination of primers that flank the *Bckdk* Flox and KO sites were used to discern the *Bckdk* Flox, WT, and KO alleles. The sequence of the flox primers were 5’-CTGCTTAAGCCCTTCCCTCT -3’ and 5’-AAGAGCACTTGCCCTTCCTT -3’. The sequence of the BCKDK KO primers were 5’-TACTGCCAGCTGGTGAGACA -3’ and 5’-GCAACACTTCCACCCAACTT -3’. The relevant product sizes were 442 bp for the WT allele, and 253 bp for the KO allele. The flox allele size is 1301 bp. The PCR was performed with the KAPA2G Fast HotStart ReadyMix (Roche, Switzerland, #KK 5609).

For *Dbt* genotyping, we selected primers which flanked the 5’ and 3’ LoxP’s to distinguish Flox, WT, and KO allele. The sequences for the forward and reverse primer are 5’-ACCGGAGCATCAGCCTAAA, and 3’-TGTGCACAAGGACATACAGG. The product sizes are 621 bp for the wild-type allele, 311 bp for the knockout allele, and 689 bp for the flox allele.

### 2.3. Developmental Assessment

A panel of developmental assessments was performed on *Bckdk*^-/-^mice and age matched littermate controls from postnatal day 1-21, unless otherwise indicated. After assessment at p1, pups were tattooed for identification throughout the study. Developmental milestones were assessed on scales or time-based assessments with a maximum cut-off time of 60 seconds. Body weight and body length from snout to tail base were measured for developmental progression. Eye opening was assessed on a scale of 0-3; 0 indicates that both eyes are closed with no slits present, 1 indicates the presence of slits but the eyes are still fully closed or one eye is partially open, 2 indicates that both eyes are partially open or one eye is fully open, and 3 indicates that both eyes are fully open. Incisor eruption was assessed as the age of full protrusion of the lower incisors through the gums. Grasping reflexes were scored on a scale of 0-2; 0 indicates no movement from either paw (no grasp), 1 indicates partial grasp or full grasp from only one paw, and 2 indicates a full grasp from both paws. Surface righting time was assessed from p4-p14 by placing mice in a supine position on a flat surface and recording the time it takes for mice to fully roll over with all four paws flat on the surface. Edge avoidance was assessed from p4-p14 by placing a mouse on a raised platform in a prone position with both the forepaws and head placed over the edge and recording the time for the mouse to retreat onto the platform with no part of the body remaining over the edge. Mice that fell from the platform could be retested a maximum of 3 times. Vertical hold time was assessed from p7-p21 by placing each mouse on top of a screen rotated 90° until vertical relative to the table. The time at which the mouse fell off the apparatus was recorded. Horizontal hold time was assessed from p10 - p21 by placing each mouse on top of a horizontal screen rotated 180° relative to the table, so that the mouse was hanging below the screen. The time to fall off the apparatus was recorded. T-bar hold time was assessed from p10-p21 by placing the forepaws of the mouse on a thin horizontal bar. Proper grip was ensured, and the time suspended on the bar was recorded. The age at maximum score or time, on scale-based or time-based assays, respectively, was also represented to illustrate developmental delay of the indicated developmental milestone. Pups that did not survive to p14 were excluded from the study.

### 2.4. µCT Analysis for Microcephaly

*Bckdk*^-/-^ mice and age matched wildtype littermate controls were collected at p21 after euthanasia with isoflurane followed by cervical dislocation and decapitation. Heads were stored in paraformaldehyde for 48 hours, then transitioned to 70% ethanol at 4°C until µCT images were obtained. Postnatal day 21 mouse skulls were scanned with a Scanco µCT35 (SCANCO Medical AG, Switzerland, Penn Center for Musculoskeletal Disorders MicroCT Imaging Core, Perelman School of Medicine, University of Pennsylvania) at a resolution of 15µm. Skulls were reconstructed with Scanco software and analysis of images was performed with 3DSlicer (V5.2.2, Slicer.org).^11^ Threshold settings were optimized to visualize only bone volume. Virtual endocasts of the skull were created to compare skull volumes by using 3DSlicer Software and the Wrap Solidify extension. Cephalometric quantifications from these images were measured from the 3D skull models including length, width, height, and endocranial volume. These measurements were inferred using previously published landmarks^12,13^. Genotype specific measurements were compared for normality and homogeneity of variance. Unpaired t-test was used for significance (p≤0.05) in the original characterization (Figure 1), 2-way ANOVA was used for BCAA treatment studies (Figure 3), and 1-way ANOVA was used for genetic rescue studies (Figure 5). Non-parametric assessment was used as needed for analyses (SPSS 26.0, IBM, Armonk, NY, USA).

**Figure. 1.**
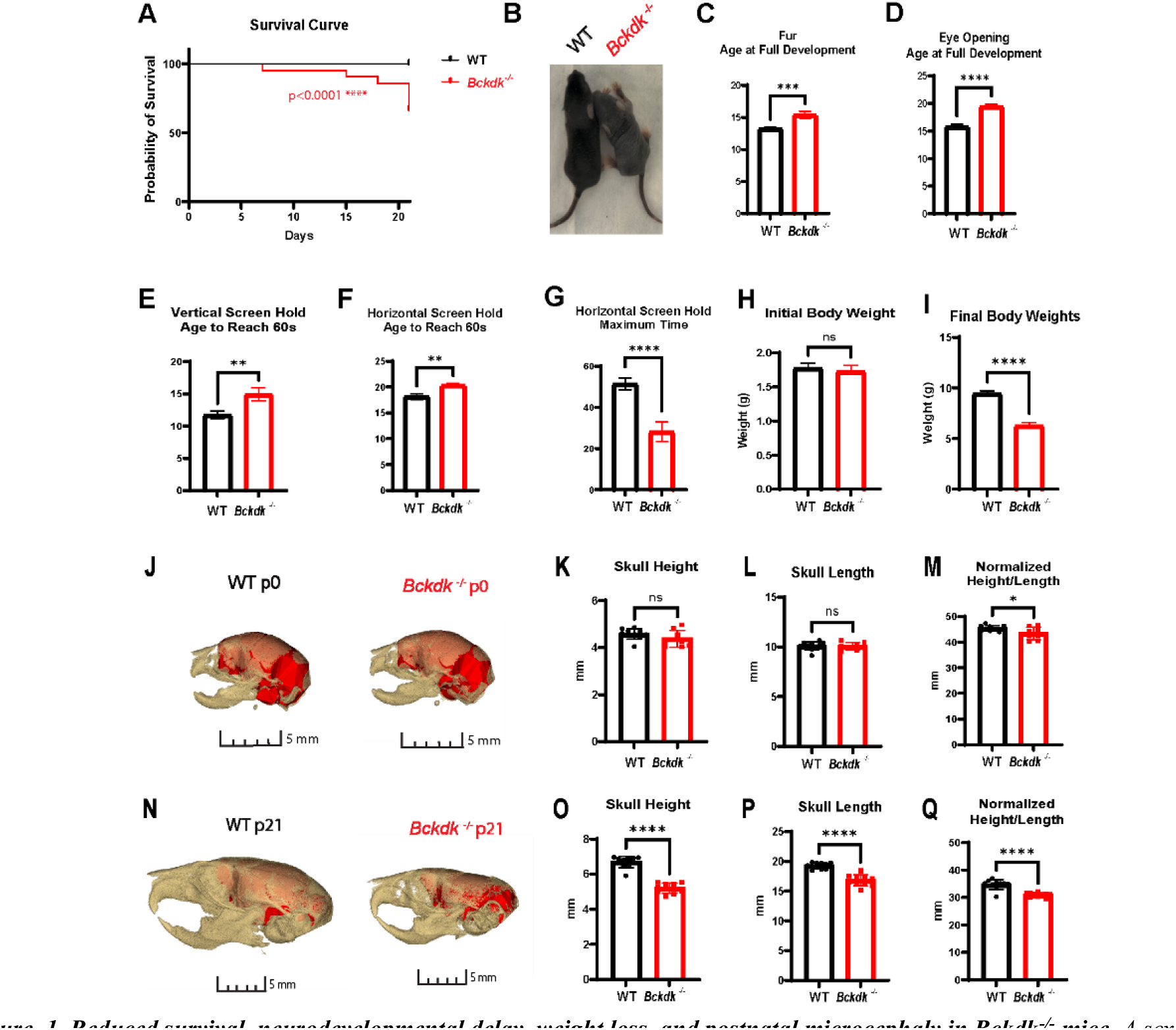
Reduced survival, neurodevelopmental delay, weight loss, and postnatal microcephaly in Bckdk^-/-^ mice. A sex balanced cohort of WT (n=27) and Bckdk^-/-^ (n-21) mice were compared for neurodevelopmental assessment. An unpaired t-test was used for statistical comparison. In all panels, data are represented as mean with s.e.m. (* p < 0.05, ** p<0.005, *** p < 0.001, **** p<0.0001, ns=non-significant). A proportion (8/21) Bckdk^-/-^ mice died before p21. (A) Fur development is delayed in Bckdk^-/-^ mice as seen by a representative image at p9. (A) A proportion (8/21) Bckdk^-/-^ mice died before p21. (B) Fur development is delayed in Bckdk^-/-^ mice as seen by a representative image at p9. (C) Full fur coat development is significantly delayed in Bckdk^-/-^ mice. (D) Age at full eye development was significantly delayed in Bckdk^-/-^ mice. (E) Delayed age to reach maximum time in Bckdk^-/-^ mice on vertical screen hold. (F) The age to reach the maximum time for horizontal screen hold is significantly delayed in Bckdk^-/-^ mice. (G) The maximum hold time is significantly reduced in Bckdk^-/-^ mice on the horizontal screen hold (max 60 seconds). (H) Similar initial body weights for Bckdk^-/-^ and control mice at p1. (I) Decrease in final body weight in Bckdk^-/-^ mice compared to WT mice at p21. (J) No changes in brain size at p0 between WT (n=10) and Bckdk^-/-^ (n=10), as seen by 3D reconstructed overlay of skull and brain from CT endocasts. (K) Minimal changes in skull height in Bckdk^-/-^ mice compared to WT mice at p0. (L) No changes in skull length in Bckdk^-/-^ mice at p0. (M) Slight decrease in a normalized metric of microcephaly (height/length*100) in Bckdk^-/-^ mice at p0. (N) Drastically smaller skull sizes in Bckdk^-/-^ (n=10) mice compared to WT (n=10) mice at p21 as seen by 3D reconstructed overlay of skull and endocast from µCT. (O) Reduced skull height in Bckdk^-/-^ mice relative to WT mice at p21. (P) Decreased skull length in Bckdk^-/-^ mice at p21 compared to WT mice. (Q) Significantly reduced skull size in Bckdk^-/-^ mice at p21 evidence of the normalized metric of microcephaly comparing skull measurements (height/length*100).

**Figure. 2.**
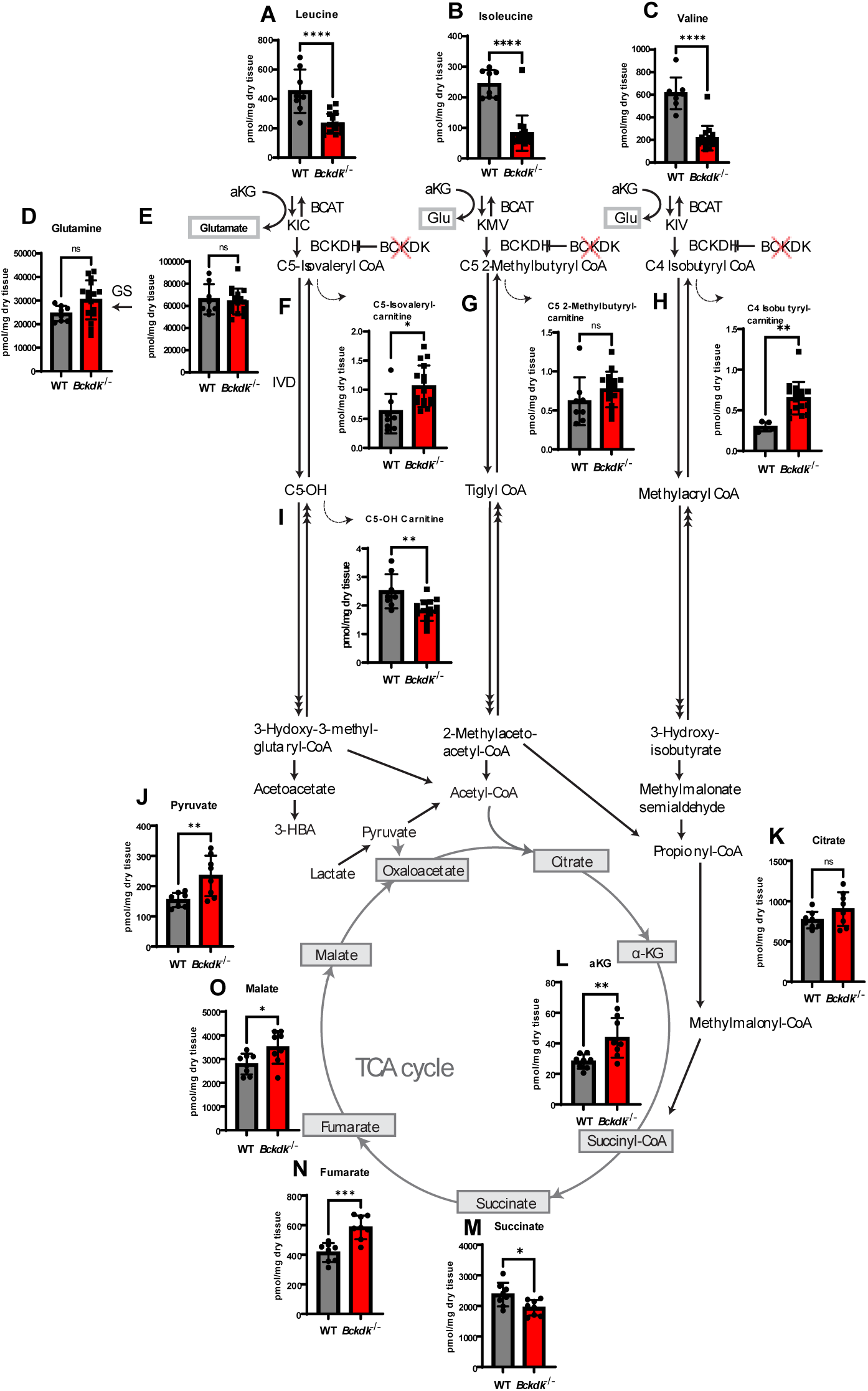
Acylcarnitine’s accumulate downstream of BCKDH in the brain of Bckdk^-/-^ mice at p21. A sex balanced cohort of WT (n=8) and Bckdk^-/-^ (n=15) mice were sent for amino acid and acylcarnitine analysis, and a subset of Bckdk^-/-^ mice (n=7) for organic acids measured by mass spectrometry. Unpaired t-tests were performed to compare the two genotypes. In all panels, data are represented as mean with s.e.m. (* p < 0.05, ** p<0.005, *** p < 0.001, **** p<0.0001, ns=non-significant). (A-C) Significant reduction of BCAA levels in Bckdk^-/-^ mice. (D-E) No changes in glutamate and glutamine levels in Bckdk^-/-^ mice. (F-H) Increased pooling of acylcarnitine intermediates one step below BCKDH in Bckdk^-/-^ mice. (I) Reduced levels of an acylcarnitine two steps below BCKDH (C5-OH) in Bckdk^-/-^ mice, specifically in the leucine degradation pathway. (J) Elevated pyruvate levels in Bckdk^-/-^ mice relative to WT levels. (K) No significant changes in citrate levels in Bckdk^-/-^ mice. (L) Increased aKG levels in Bckdk^-/-^ mice. (M) Decreased succinate levels in Bckdk^-/-^ mice. (N) Significant increase in fumarate levels in Bckdk^-/-^ mice. (O) Increased malate levels in Bckdk^-/-^ mice.

**Figure. 3.**
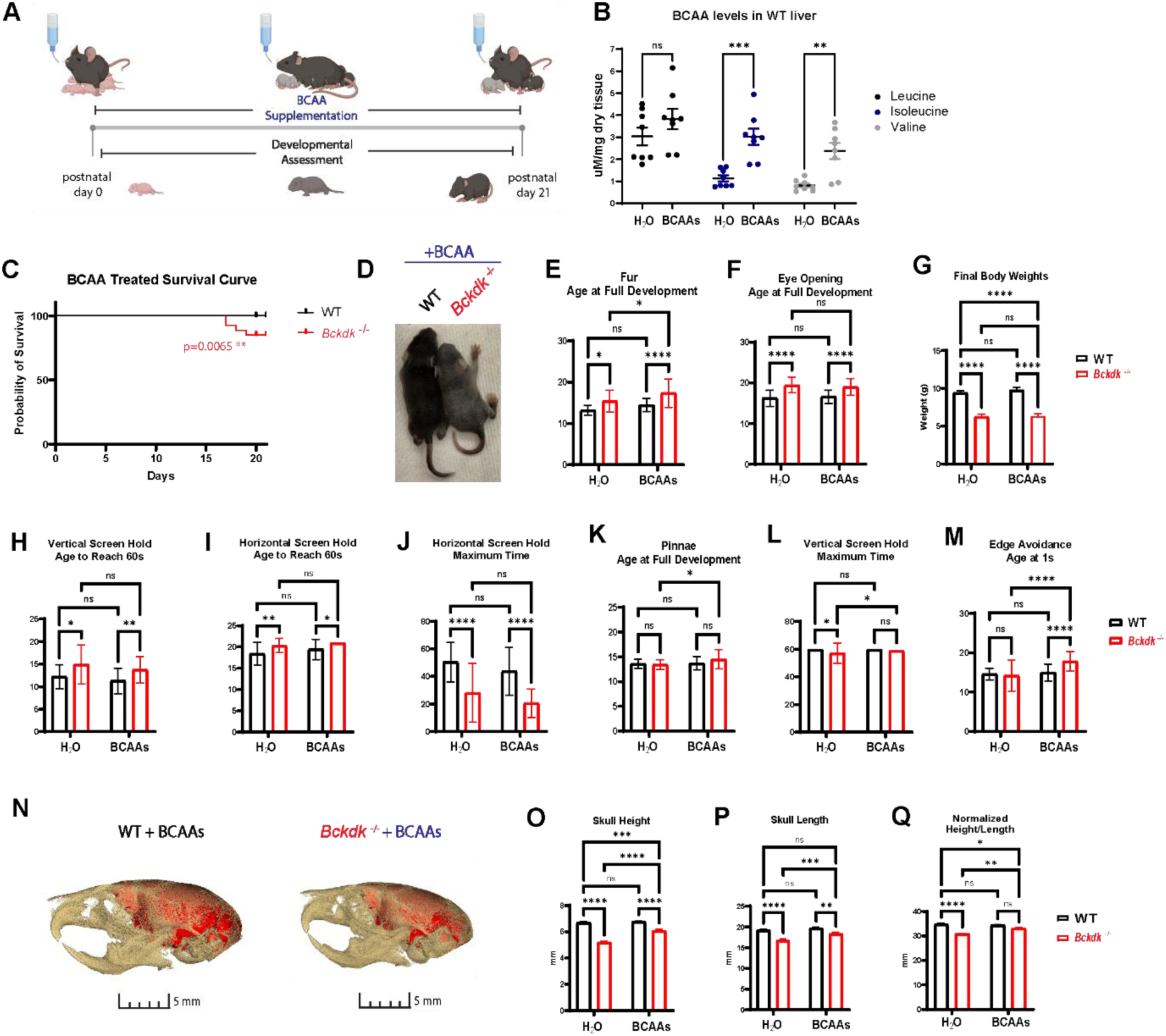
BCAA supplementation did not significantly impact survival, neurodevelopmental delay, weight loss, or microcephaly in Bckdk^-/-^ mice. A sex balanced cohort of WT (n=39) and Bckdk^-/-^ (n=28) mice were compared for all neurodevelopmental assessments. A 2-way ANOVA with multiple comparisons was used for all statistical analyses. In all panels, data are represented as mean with s.e.m. (* p < 0.05, ** p<0.005, *** p < 0.001, **** p<0.0001, ns=non-significant). Experimental design of BCAA administration through dams through lactation, with simultaneous neurodevelopmental assessment of treated pups. (A) Significant elevation of BCAAs in the livers of BCAA treated WT mice (n=8). (B) Reduced survival (4/28; 14%) of BCAA treated Bckdk^-/-^ mice prior to p21. (C) Delayed fur development of BCAA treated Bckdk^-/-^ mice as seen by representative images at p9. (D) Delayed age of full coat development of BCAA treated Bckdk^-/-^ mice. (E) Delayed age of full eye opening in BCAA treated Bckdk^-/-^ mice. (F) Significantly reduced final body weight in BCAA treated Bckdk^-/-^ mice at p21. Final body weight was taken from mice collected for experiments (WT(n=8), BCKDK^-/-^(n=16), BCAA treated WT (n=9), BCAA treated Bckdk^-/-^ (n=8)). (G) Delayed age to reach the maximum hold time for vertical screen in BCAA treated Bckdk^-/-^ mice. (H) Delayed age to reach the maximum hold time for horizontal screen in BCAA treated Bckdk^-/-^ mice. (I) Decreased maximum hold time on horizontal screen hold in BCAA treated Bckdk^-/-^ mice. (J) Further delayed age to reach full pinnae development in BCAA treated Bckdk^-/-^ mice compared to untreated Bckdk^-/-^ mice. (K) Further increased maximum time on vertical screen hold in BCAA treated Bckdk^-/-^ mice compared to untreated Bckdk^-/-^ mice. (L) Further worsening of edge avoidance in BCAA treated Bckdk^-/-^ mice compared to untreated Bckdk^-/-^ mice. (M) Smaller skull sizes in BCAA treated Bckdk^-/-^ (n=10) mice compared to BCAA treated WT (n=10) mice at p21 as seen by 3D reconstructed overlay of skull and endocast from µCT. (N) Reduced skull height in BCAA treated Bckdk^-/-^ mice (n=10 per group). (O) Decreased skull length in BCAA treated Bckdk^-/-^ mice (n=10 per group). (P) Reduced skull size in BCAA treated Bckdk^-/-^ mice at p21 evidence of the normalized metric of microcephaly comparing skull measurements.

### 2.5. Amino acid levels analysis

The concentration of amino acids was determined by HPLC, Agilent Technologies 1260 Infinity II system, utilizing precolumn derivatization with *o*-phthalaldehyde (OPA) as described^30^. Briefly, in each HPLC chromatogram we determined the profile of a-amino acids in the plasma following precolumn derivatization with OPA. The level of each amino acid was determined relative to a known concentration of internal standard, e-amino-caproic acid (EACA), added to plasma sample. For the BCAA levels measured using the Branched Chain Amino Acid Assay Kit (Cell Biolabs, Inc. San Diego, CA, #MET-5056), tissue homogenates were prepared in PBS, sonicated, and centrifuged to collect the supernatants for measurements. A standard curve was prepared using the L-leucine standard included in the kit. The sample supernatants were then mixed with assay buffer, WST-1 Reagent, NAD+, and leucine dehydrogenase according to the manufacturer’s instructions. After 5 – 10 minutes incubation, a plate reader read the absorbance at 450 nm. Reagents were aliquoted to avoid multiple freeze/thaw cycles, and the leucine dehydrogenase solution was prepared fresh before use. BCAA levels were normalized to protein concentrations as measured by a BCA protein assay (ThermoFisher Scientific, #23227).

Liver tissue from BCAA supplemented animals were snap frozen in liquid nitrogen. Frozen tissue was homogenized, and 30 mg was weighed out. Tissue was mixed with 1 mL 40:40:20 methanol:acetonitrile:water (extraction solvent) and homogenized with a benchtop lyser (SciLogex OS20-S) at 1600 mhZ for approximately 15 sec. U-13C-propionate was spiked in to the extraction solvent as an internal standard. Samples were then immediately centrifuged at 13.3g for 20 mins at 4C. Supernatant was saved and dried down in a speedvac (ThermoFisher Savant SpeedVac SPD130DLX) overnight at room temp. Samples were resuspended in 50 uL 60:40 methanol:water and spun down again at 13.3g for 20 mins at 4C. Supernatant was collected for LC-MS analysis.

A quadrupole-orbitrap mass spectrometer (Q Exactive, Thermo Fisher Scientific, San Jose, CA) operating in negative ion mode was coupled to hydrophilic interaction chromatography on a Vanquish UHPLC System (Thermo Fisher Scientific) via electrospray ionization and used to scan from m/z 65-425 at 1 Hz and 140,000 resolution. LC separation was on a ACQUITY Premier BEH Amide VanGuard FIT Column, 1.7 µm, 2.1 mm X 100 mm (Water, Milfod, MA) using a gradient of solvent A (20 mM ammonium acetate, 20 mM Ammonium hydroxide in 95:5 water:acetonitrile, pH 9.45) and solvent B (acetonitrile). Flow rate was 300 µL/min. The LC gradient was: 0 min, 95% B; 9 min, 40%; 11 min, 40%; 11.1 min, 95%; 20 min, 95%. Autosampler was set at 4°C and injection volume was 3 µL. LC-MS was used to measure BCAAs and BCKAs. A standard curve for BCAAs/BCKAs was run in parallel to allow for quantitation. Data analysis was done using El-MAVEN software^14^.

### 2.6. BCAA Supplementation

BCAA powder was ordered at a premixed 2:1:1 ratio of Leu:Iso:Val (BCAA 5000 Natural Powder, Nutrabio, Inc., Middlesex, NJ). A previous publication verified this biochemical ratio with assistance from an independent 3^rd^ party vendor (ALS Global, Salt Lake City, UT).^15^ A 2% percent BCAA solution (2g/100mL) was introduced via drinking water to dams with a new litter at age p0/p1 and maintained until p21. Bottles were weighed alternate days to monitor intake and the solution was refreshed weekly. IACUC approval was received for BCAA supplementation.

### 2.7. Realtime qPCR

RNA extraction, cDNA synthesis, and real-time qPCR were performed as described^31^. Briefly, RNA was extracted and purified using a combination of TRIzol reagent and QIAGEN RNeasy columns (QIAGEN, #74004); reversed transcribed into cDNA using the qScript cDNA SuperMix (Quantabio, Beverly, MA, #95047). The cDNA was then diluted and used as the template for real-time PCR analysis using the Power up SYBR Green PCR Master Mix (ThermoFisher Scientific, # A25779) on the QuantStudio3 Real-Time PCR System (ThermoFisher Scientific). The list of primers and sequences used is found in Supplementary Table 1.

### 2.8. Western Blot

Whole brains were homogenized in 1X RIPA buffer followed by BCA assay (ThermoFisher Scientific, #23227). Equivalent protein quantities for all genotypes were loaded onto a Nupage 4-12% Bis-Tris Protein Gels (ThermoFisher Scientific) for 1 hour. Subsequently, resolved proteins were transferred to a nitrocellulose membrane for 1.5 hours (ThermoFisher Scientific). Membranes were blocked with 5% bovine serum albumin in tris-buffered saline with 0.1% tween (TBS-T) for 1 hour, followed by incubation of primary antibody dilutions in blocking solution overnight at 4°C. The following primary antibodies were used for Western blot including BCKDK, pBCKDH-E1a, BCKDH-E1a, and GAPDH. Additional antibody information can be found in Supplementary Table 2. Membranes were washed with TBS-T, incubated with HRP-conjugated secondary antibodies (1:10,000) diluted in blocking solution for 1 hour, and washed with TBS-T prior to imaging. SuperSignal™ West Pico PLUS reagents were used (ThermoFisher Scientific) to image relative protein amounts using a ChemiDoc Gel Imaging System (BioRad, Hercules, CA). After acquisition, protein levels were quantified using Image J software’s band analysis feature (NIH).

### 2.9. Statistical analysis

Data are presented as mean ± SEM. Statistical analysis was performed using Prism8 software (GraphPad, San Diego, CA). The respective statistical test for each experiment is indicated in the figure legends. Regardless of statistical test performed, statistical significance was annotated with asterisks within the figures as follows for all tests and results, unless otherwise stated (* p <0.05, ** p <0.005, *** p <0.0005, **** p <0.0001).

## 3. Results

### 3.1. *Bckdk^-/-^* mice have reduced survival, developmental delay, and postnatal microcephaly

To determine if a previously published *Bckdk*^-/-^ model had clinically relevant neurological phenotypes of BCKDK deficiency, a panel of neurodevelopmental assessments and micro computed tomography (µCT) analysis for microcephaly was performed^10^. We validated molecular loss of BCKDK by PCR and western blot in addition to the phosphorylation site of BCKDH-E1a subunit indicating functional loss of BCKDK (Supplemental Figure 1A-D). A subset (38%) of *Bckdk*^-/-^ mice died prior to weaning at postnatal day 21 (p21) (Figure 1A).

Prior to p21, neurodevelopmental assessments revealed that ectodermal-derived tissues have delayed development including delayed fur development (Figure 1B-C) and eye opening (Figure 1D). Motor development was also reduced with delayed age to reach the maximum screen hold times in both a vertical and horizontal assay (Figure 1E-G, Supplemental Figure 1J). The initial weight of *Bckdk*^-/-^ mice was not significantly different from WT controls (Figure 1H). *Bckdk*^-/-^ mice gained weight during the first two weeks of life, but then started losing weight in the third week, indicative of developmental regression (Supplemental Figure 1E). Final weight at p21 was significantly reduced in *Bckdk*^-/-^mice (Figure 1I). Additional metrics assessed were not significantly altered including mouse length, grasping ability, pinnae development, incisor protrusion, maximum time for T-bar suspension, surface righting, and edge avoidance (Supplemental Fig 1F-I,K-M). Microcephaly was assessed by µCT at p0, and there was an initiation of a subtle reduction in skull height to skull length at this early age (Figure J-M). µCT derived endocasts of skulls at p0 revealed no differences in endocranial volume, normalized to skull length, at this early age (Supplemental Figure 1N-O). However, by p21 *Bckdk*^-/-^ mice had significantly reduced skull height to skull length, revealing a flattened skull characteristic of postnatal microcephaly and suggesting a delay in cortical development (Figure 1N-Q)^16^. Similarly, *Bckdk*^-/-^ mice had noticeably smaller brains seen by 3D endocasts and significantly reduced endocranial volumes at p21, revealing that microcephaly arises postnatally and that brain size is affected (Supplemental Figure 1P-Q).

Quantification of physical brain size revealed reduced brain weight recapitulating µCT results at p21 (Supplemental Figure 1R-S). Therefore, this *Bckdk*^-/-^ mouse effectively models clinically relevant neurological phenotypes of human disease including neurodevelopmental delay and postnatal microcephaly.

### 3.2. Acylcarnitine and TCA cycle intermediate pooling in *Bckdk^-/-^* brains

Previous literature revealed depletion of BCAAs in human patients and a decrease in downstream adenosine triphosphate (ATP) levels in BCKDK deficient patient-derived cells.^5,7–10,17,18^ Given the unique role of BCAAs in the brain, we sought to elucidate how BCKDK deficiency impacts BCAA catabolism and downstream TCA cycle intermediate levels in the brain. We hypothesized that there would be increased levels of intermediates derived from BCAA nitrogen donation (glutamate, glutamine) and carbon donation (acylcarnitine species, TCA cycle intermediates) due to loss of BCKDK. Brain BCAA levels were significantly reduced at p21, as has previously been shown at a similar timepoint in another *Bckdk*^-/-^ mouse model (Fig2A-C)^17^. Glutamate and glutamine levels were unchanged from WT levels at p21, implying that either insignificant changes in nitrogen donation or compensation after increased nitrogen donation to regulate levels in the brain (Figure 2D-E). Either possibility suggests it is an unlikely mechanism of biochemical dysregulation. Acylcarnitine conjugates of downstream intermediates below the BCKDH step accumulated, supporting that there is increased carbon donation to direct downstream intermediates (Figure 2F-H). Interestingly, the leucine-derived intermediate C5-OH carnitine is depleted, two steps below the BCKDH step, suggesting a blockade in catabolism and pooling of upstream intermediates (Figure 2I)^19^.

To bridge the gap between pooling BCAA acylcarnitine intermediates and their downstream impact on downstream metabolism in the brain, we investigated if organic acids, particularly TCA-cycle intermediates, were altered. 3-HBA organic acid levels were elevated in *Bckdk*^-/-^ mice (Supplemental Figure 2A). Pyruvate levels, derived from both glucose and BCAA breakdown, were elevated while lactate levels were similar between groups, suggesting that cytosolic NAD/NADH ratios may be increased (Figure 2J, Supplemental Figure 2B). All TCA cycle intermediate levels increase, except succinate, implying that there is increased anaplerosis in *Bckdk*^-/-^ mice (Figure J-O). These results are consistent with higher flux through the anaplerotic BCAA catabolic pathways.

Interestingly, we also found a blockade in other amino acid degradation pathways that have been previously implicated in neurologic disorders, specifically lysine and glycine degradation. Lysine and tryptophan levels accumulated in *Bckdk*^-/-^ mice (Supplemental Figure 2C-D). Accumulation of lysine degradation intermediates have previously been associated with pyridoxine-dependent epilepsy^20–22^. Therefore, we looked at downstream C5-DC levels, which are also significantly depleted implying that reduced lysine degradation (Supplemental Figure 2E). In addition, glycine levels were significantly elevated at p21 (Supplemental Figure F). Glycine has been associated with glycine encephalopathy, a severe seizure disorder^23^. Clinical biomarkers of BCKDK deficiency, tyrosine, phenylalanine, and alanine, were significantly and trend increased in *Bckdk*^-/-^ mice (Supplemental Figure 1G-I). Subtle abnormalities in additional amino acid levels were also observed (Supplemental Figure 1J-R).

### 3.3. BCAA repletion does not rescue survival, developmental delay, or microcephaly

To determine if BCAA repletion could impact clinically relevant neurological phenotypes in our *Bckdk*^-/-^ model, we performed the same panel of neurodevelopmental assessments and µCT evaluation for microcephaly in BCAA treated mice and compared them to our untreated mice^10^. BCAAs were diluted in reverse osmosed water to a supersaturated 2% solution, as previously described^15^. BCAAs were administered through lactation to treat pups continuously throughout the day to maximize BCAA repletion from p0-p21 (Figure 3A).

BCAAs did not significantly impact overall water intake and daily dosage was adjusted for litter size by maternal intake (Supplemental Figure 3A-B). Through this administration route, BCAAs were significantly increased in the liver of WT pups (Figure 3B). BCAA treated *Bckdk*^-/-^ mice still had 15% reduced survival, revealing peripherally administered BCAAs did not fully rescue survival deficits (Figure 3C). Developmental delay of ectodermal-derived tissues persisted in BCAA treated *Bckdk*^-/-^ pups including delayed fur development (Figure 3D-E), and eye opening (Figure 3F). BCAA treated *Bckdk*^-/-^ mice initially gained weight similarly to control littermates, but also developmentally regressed around week 2 of life (Supplemental Figure 3C). Final body weight was not significantly altered by BCAA administration as compared to untreated *Bckdk*^-/-^ mice (Figure 3G). Motor development was also delayed despite BCAA treatment, with reduced age to reach the maximum time in the horizontal screen hold and reduced screen hold times in both the vertical and horizontal assay (Figure 3H-K). BCAA administration worsened maximum time on the vertical screen hold, pinnae development, and edge avoidance in BCAA treated versus untreated *Bckdk*^-/-^ mice (Figure 3K-M). Furthermore, differences in edge avoidance, grasping, and maximum T-bar suspension time were new defects elicited by BCAA administration to *Bckdk*^-/-^ mice (Figure 3M, Supplemental Figure 3D-E). Similar to our initial characterization, body length, incisor protrusion, and surface righting did not differ between groups (Supplemental Figure 3F-H).

Microcephaly was not drastically improved by BCAA treatment, as skull height and lengths were still reduced (Figure 3N-Q)^16^. BCAA treatment did increase brain weight (Supplemental Figure 3K-L), which recapitulates what was seen in the first *Bckdk*^-/-^ mouse model supplemented with a high protein diet^8^. However, BCAA treatment was not sufficient to normalize endocranial volumes (Supplemental Figure I-J). Together, these data indicate that postnatal peripheral BCAA repletion is not sufficient alone to rescue neurodevelopmental delay in *Bckdk*^-/-^ mice and can even worsen some neurological phenotypes.

### 3.4. BCAA repletion reduced catabolic pathway pooling but not downstream TCA cycle changes in the brain of *Bckdk*^-/-^ mice

Although BCAA repletion did not significantly rescue neurodevelopmental delay in mice, it remained unclear if BCAA repletion would be sufficient to prevent pooling of pathway intermediates and normalize downstream TCA cycle intermediate levels. Peripheral BCAA supplementation through lactation did not drastically elevate BCAA levels in the brain despite elevated levels in the liver (Figure 4A-C), revealing a limitation of peripheral BCAA intervention also seen in a lack of restoration of BCAA levels in the central nervous system in human patients^7^. Again, glutamate and glutamine levels were not altered by this intervention (Figure 4D-E). BCAA supplementation did reduce the amount of pooling of intermediates directly downstream of the BCKDH step, as measured through acylcarnitine analysis (Figure 4F-H). The intermediate two steps down from BCKDH, C5-OH carnitine, was further depleted in *Bckdk*^-/-^ mice with BCAA supplementation (Figure 4I).

**Figure. 4.**
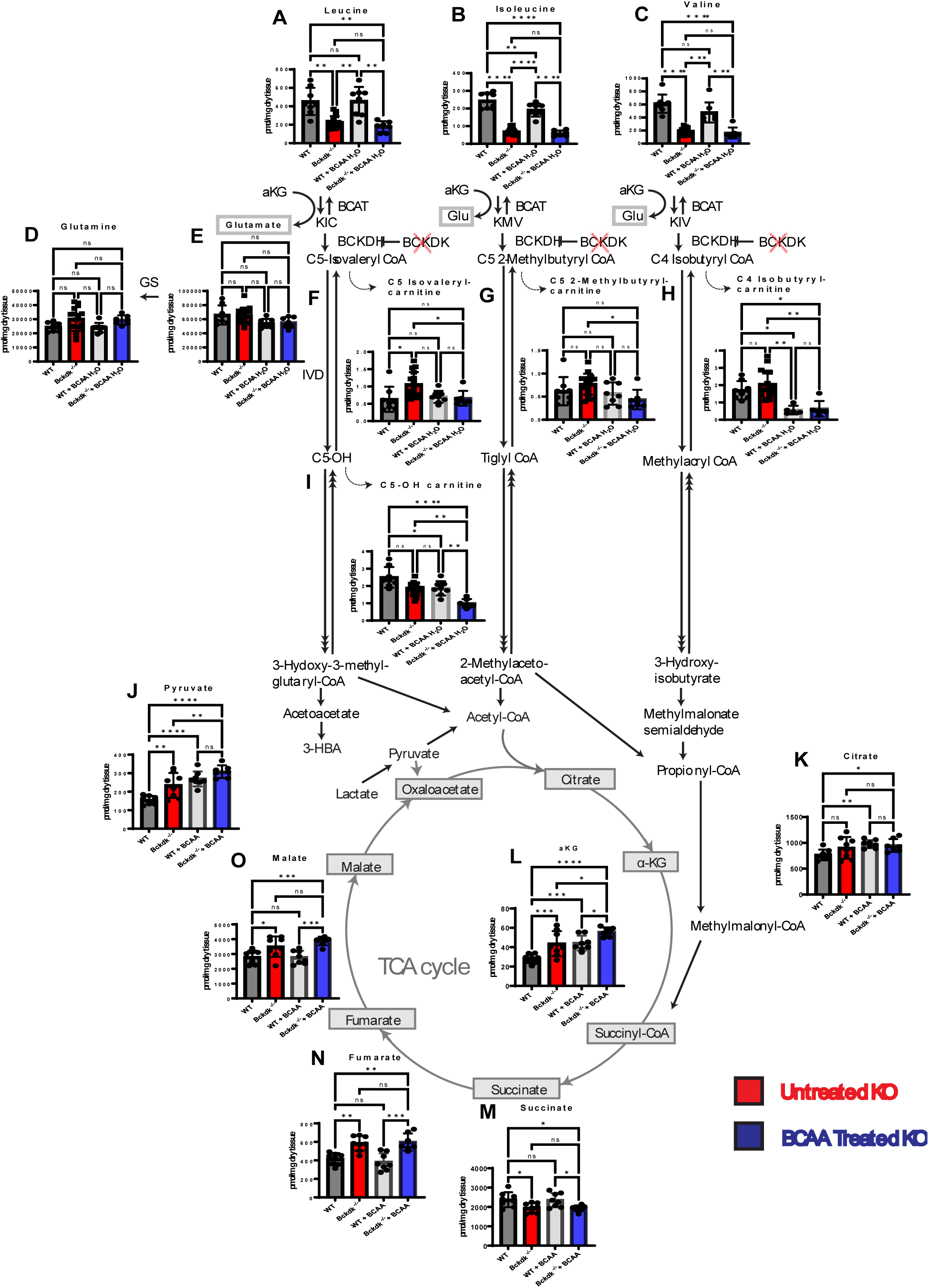
Amino Acids and Acylcarnitine levels in the brain of WT and Bckdk^-/-^ mice at p21. A sex balanced cohort of BCAA treated WT (n=8) and Bckdk^-/-^ (n=7) mice were compared to untreated WT (n=8) and Bckdk^-/-^ (n=15) data from figure 2. Brain amino acid, acylcarnitine, and organic acid levels were measured by mass spectrometry. 2-way ANOVAs were performed to compare all four groups for genotype and BCAA intervention differences. In all panels, data are represented as mean with s.e.m. (* p < 0.05, ** p<0.005, *** p < 0.001, **** p<0.0001, ns=non-significant). (A-C) BCAA levels are significantly reduced in BCAA treated Bckdk^-/-^ mice at p21. (D-E) No differences in glutamate and glutamine in BCAA treated Bckdk^-/-^ mice. (F-H) Reduced pooling of acylcarnitine intermediates one step below BCKDH in BCAA treated Bckdk^-/-^ mice compared to untreated Bckdk^-/-^ mice. (I) Further reduced levels of acylcarnitine species two steps below BCKDH (C5-OH) in BCAA treated BCKDK^-/-^ mice relative to untreated Bckdk^-/-^ mice. (J) Further elevated pyruvate levels in BCAA treated Bckdk^-/-^ mice compared to untreated Bckdk^-/-^ mice and untreated WT mice. (K) Increase in citrate levels in BCAA treated Bckdk^-/-^ mice relative to untreated WT mice. (L) Further increased aKG levels in BCAA treated Bckdk^-/-^ mice compared to WT levels and untreated Bckdk^-/-^ levels. (M) Similarly decreased succinate levels in BCAA treated Bckdk^-/-^ mice compared to untreated Bckdk^-/-^ mice. (N) Similar increase in fumarate levels in BCAA treated Bckdk^-/-^ mice relative to Bckdk^-/-^ mice. (O) Further increase in malate levels in BCAA treated Bckdk^-/-^ mice compared to untreated Bckdk^-/-^ mice, both relative to WT levels.

Since BCAA supplementation corrected levels of upstream intermediates, we investigated if BCAA supplementation also normalized organic acid levels, particularly downstream intermediates of the TCA cycle. 3-HBA levels were elevated upon BCAA supplementation in *Bckdk*^-/-^ mice (Supplemental Figure 4A). Pyruvate levels were further elevated by BCAA supplementation (Figure 4J), while lactate levels remained unchanged (Supplemental Figure 4B). Citrate levels were increased compared to WT untreated controls (Figure 4K).

Interestingly, aKG levels were further increased on BCAA supplementation in *Bckdk*^-/-^ mice (Figure 3L). Inversely, succinate levels were still depleted in *Bckdk*^-/-^ mice even in the presence of BCAAs (Figure 4M). Fumarate and malate levels were elevated with similar levels compared to untreated *Bckdk*^-/-^ mice (Figure 4N-O). The pooling of citrate and aKG is consistent with increased catabolism upon BCAA supplementation in *Bckdk*^-/-^ mice, although this is not reflected in the fumarate-malate span of the TCA cycle. Together, these data show that BCAA supplementation has identifiable effects on levels of various brain metabolic intermediates, but it alone does not normalize carbon donation to downstream intermediates nor restore TCA cycle changes due to loss of BCKDK.

Lysine accumulation in BCAA treated *Bckdk*^-/-^ mice remained increased while lysine degradation remained restricted as evidence by reduced C5-DC carnitine (Supplemental Figure 4C-E). Similarly, glycine levels remained elevated in untreated *Bckdk*^-/-^ mice despite BCAA treatment (Supplemental Figure 4F). Additional amino acid markers of BCKDK deficiency remained elevated in BCAA treated *Bckdk*^-/-^ mice compared to BCAA treated WT mice (Supplemental Figure 4G-I). Multiple amino acids were not impacted by BCAA supplementation including arginine, histidine, and aspartate (Supplemental Figure 4J-K, P). Specific amino acids, proline and asparagine, were similarly elevated in *Bckdk*^-/-^ mice with or without BCAA supplementation (Supplemental Figure 4N-O). BCAA supplementation further exacerbated differences in amino acid levels particularly for serine between *Bckdk*^-/-^ and WT treated mice (Supplemental Figure 4M,O,Q-R). Together this data suggests that peripheral repletion of BCAAs does not restore all metabolic changes in the brain nor neurodevelopmental delay phenotypes associated with BCKDK deficiency, implying that an alternative mechanism of disease pathology besides BCAA depletion alone.

### 3.5. DBT haploinsufficiency rescues neurodevelopmental delay in *Bckdk^-/-^* mice

An alternative hypothesis to low levels of BCAAs being the cause of neurological defects in *Bckdk^-/-^* mice is the hypothesis that high flux of BCAA catabolism is in fact causal. To test this hypothesis, we reduced BCAA catabolism by haploinsufficiency of *Dbt*, encoding the critical E2 subunit of the BCKDH complex. We crossed *Bckdk^+/-^* mice with a novel *Dbt^+/-^* mouse model to obtain *Bckdk ^+/-^ Dbt ^+/-^* mice for breeding. These mice were further crossed to obtain “genetic rescue” mice (*Bckdk^-/-^Dbt^+/-^)* and littermate controls for neurodevelopmental assessment and metabolic analysis (Figure 5A). *Dbt* haploinsufficiency was detected by PCR, and DBT mRNA levels in genetic rescue mice were decreased 32% relative to normalized WT levels in the brain (Supplemental Figure 5A-B, Figure 5B-C).

**Figure. 5.**
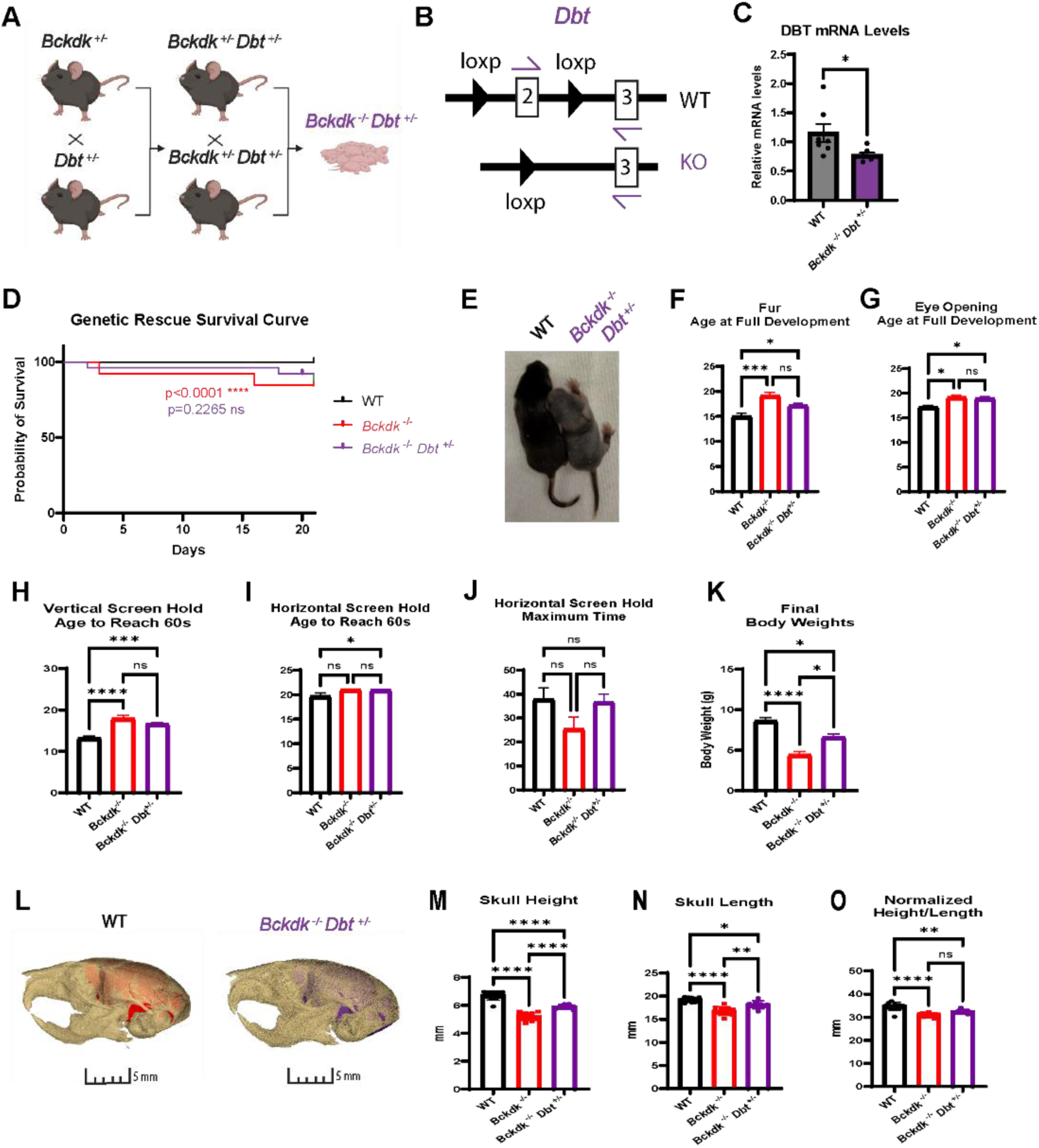
Higher survival, less severe neurodevelopmental delay, increased weight gain, and partial restoration of microcephaly in Bckdk^-/-^ Dbt^+/-^ mice. A sex balanced cohort of WT (n=12), Bckdk^-/-^ (n=13), and Bckdk^-/-^Dbt^+/-^ (n=26) mice were compared for all neurodevelopmental assessments. A one-way ANOVAs with multiple comparisons was used for comparisons between the three groups. In all panels, data are represented as mean with s.e.m. (* p < 0.05, ** p<0.005, *** p < 0.001, **** p<0.0001, ns=non-significant). (A) Breeding scheme to obtain Dbt haploinsufficient Bckdk^-/-^ mice. Bckdk^+/-^ and Dbt^+/-^ mice were crossed to obtain Bckdk^+/-^Dbt^+/-^. These mice were further crossed to obtain Bckdk^-/-^Dbt^+/-^ mice. (B) Design of qRT-PCR for Dbt KO and WT allele to measure mRNA transcript levels. (C) Decreased relative DBT transcript levels in Bckdk^-/-^Dbt^+/-^ (n=7) mice compared to WT (n=7) mice measured by qRT-PCR and normalized to Actin levels. (D) Higher survival rates in Bckdk^-/-^Dbt^+/-^ (3/25;12%) mice compared to littermate matched Bckdk^-/-^ mice (3/13;23%). (E) Delayed fur development in Bckdk^-/-^Dbt^+/-^ as seen by representative images at p10. (F) Partial rescue of delayed age of full coat development of Bckdk^-/-^Dbt^+/-^ mice relative to Bckdk^-/-^ mice, both compared to WT mice. (G) Similarly delayed age of full eye opening in Bckdk^-/-^Dbt^+/-^ mice compared to Bckdk^-/-^ mice. (H) Similarly delayed age to reach the maximum hold time for vertical screen in Bckdk^-/-^Dbt^+/-^ mice relative to Bckdk^-/-^ mice. (I) Later age to reach the maximum hold time for horizontal screen in Bckdk^-/-^Dbt^+/-^ mice compared to WT mice. (J) Similar maximum time on horizontal screen hold in Bckdk^-/-^Dbt^+/-^ mice relative to WT mice. (K) Final body weights are partially rescued in Bckdk^-/-^Dbt^+/-^ compared to WT and Bckdk^-/-^ matched controls. (L) Partial rescue of skull size in Bckdk^-/-^Dbt^+/-^ (n=10) mice compared to WT (n=10) mice at p21 as seen by 3D reconstructed overlay of skull and endocasts from µCT. (M) Partially restored increase in skull height in Bckdk^-/-^Dbt^+/-^ mice (n=10 per group) compared to Bckdk^-/-^ mice. (N) Partial rescue of skull length in Bckdk^-/-^Dbt^+/-^ mice (n=10 per group) compared to WT mice. (O) Partial restoration of skull size in Bckdk^-/-^Dbt^+/-^ mice at p21 compared to WT mice evidence of the normalized metric of microcephaly comparing skull measurements.

DBT haploinsufficiency slightly increased *Bckdk*^-/-^ mice survival with 12% death prior to our endpoint at 21 (Figure 5D). Prior to this endpoint, neurodevelopmental assessment revealed a trend rescue of fur development (Figure 5E-F), but no change in the age at full eye opening (Figure 5G). Motor development was partially rescued with slightly earlier ages to reach the maximum time of vertical screen hold (Figure 5H). Although the age to be able to reach the maximum time on the horizontal screen was delayed compared to WT mice, their average time on the horizontal screen was more similar to WT mice than *Bckdk*^-/-^ mice indicative of motor improvement (Figure 5I-J). Genetic rescue mice tracked with WT mice in their weight gain throughout the 3-week assessment (Supplemental Figure 5C). Additionally, final body weights at p21 were partially rescued (Figure 5K). Genetic rescue mice had similar lengths, grasping, pinnae development, incisor protrusion, max vertical screen hold time, maximum suspension time on T-bar, surface righting, and edge avoidance compared to both WT and *Bckdk^-/-^* mice (Supplemental Figure 5D-K). Microcephaly analyses revealed highly significant partial rescue of skull length (Figure 5L-O)^16^. Estimation of brain size through µCT analysis indicated higher endocranial volume relative to *Bckdk*^-/-^ mice (Supplemental Figure 5L-M). Partial rescue of brain weight recapitulated CT studies with a trend towards increased brain weight relative to *Bckdk*^-/-^ mice (Supplemental Figure 5N-O). These data thus support the use of *Dbt* inhibition as a novel therapeutic target, which partially rescues neurodevelopmental delay and microcephaly in *Bckdk*^-/-^ mice.

### 3.6. DBT haploinsufficiency reduces BCAA catabolite pooling and partially rescues TCA cycle changes in the brain of *Bckdk*^-/-^ mice

Next, we evaluated if *Dbt* haploinsufficiency could improve metabolic markers of BCKDK deficiency in the brain. DBT haploinsufficiency in genetic rescue mice did not significantly increase BCAA levels in the brain (Figure 6A-C), further supporting that BCAA depletion alone is not the sole cause of disease pathology. However, DBT haploinsufficiency was sufficient to reduce pooling of intermediates one step below BCKDH, partially normalizing the BCAA catabolic pathway (Figure 6F-I). Glutamate and glutamine levels were not significantly changed similar to control groups (Figure 6D-E), again supporting that excess glutamate is an unlikely cause of disease pathology in BCKDK deficiency.

**Figure. 6.**
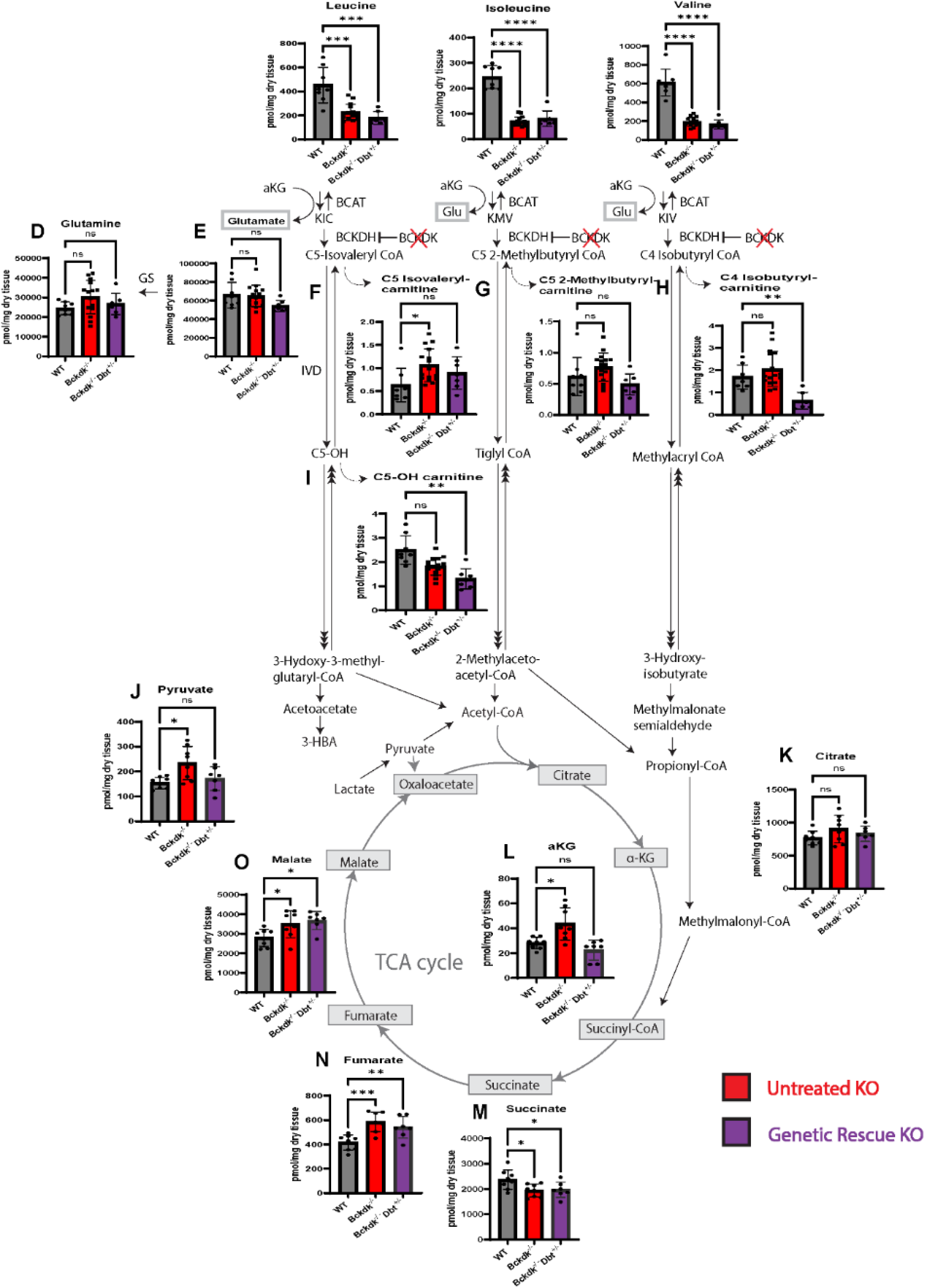
Amino Acids and Acylcarnitine levels in the brain of WT and Bckdk^-/-^Dbt^+/-^ mice at p21. A sex balanced cohort of Bckdk^-/-^Dbt^+/-^ (n=7) mice were compared to untreated WT (n=8) and Bckdk^-/-^ (n=15) metabolic data from the original characterization shown in Figure 2. Amino acid, acylcarnitine, and organic acids levels were measured by mass spectrometry. One-way ANOVA with multiple comparisons was used to compare genotypes. In all panels, data are represented as mean with s.e.m. (* p < 0.05, ** p<0.005, *** p < 0.001, **** p<0.0001, ns=non-significant). (A-C) Significant reduction of BCAA levels in Bckdk^-/-^Dbt^+/-^ mice at p21. (D-E) No changes in glutamate and glutamine in all groups at p21. (F-H) Reduced pooling of acylcarnitine species one step below BCKDH are relatively similar in Bckdk^-/-^Dbt^+/-^ mice compared to WT mice, and C4-Isobutyryl carnitine was even further reduced. (I) Further reduced levels of acylcarnitine species two steps below BCKDH (C5-OH) in Bckdk^-/-^Dbt^+/-^ mice relative to WT mice. (J) Normalization of pyruvate levels back to WT levels in Bckdk^-/-^Dbt^+/-^ mice. (K) Citrate levels were similar in all groups. (L) aKG levels resolved in Bckdk^-/-^Dbt^+/-^ mice back to WT levels. (M) Succinate levels were similarly decreased in Bckdk^-/-^Dbt^+/-^ mice compared to WT mice. (N) Fumarate levels remained elevated in Bckdk^-/-^Dbt^+/-^ mice relative to WT, although to a lesser extent than Bckdk^-/-^ levels. (O) Malate levels were also elevated in Bckdk^-/-^Dbt^+/-^ mice compared to WT mice, having similar levels to Bckdk^-/-^ mice.

Lastly, we investigated if DBT haploinsufficiency in BCKDK deficiency restored organic acid including TCA cycle intermediate changes. Pyruvate levels were restored back to WT levels (Figure 6J). Citrate levels remained unchanged (Figure 6K). aKG levels resolved back to WT levels in genetic rescue mice (Figure 6L), suggesting there is partial correction of the TCA cycle. However, downstream changes in succinate, fumarate, and malate were unchanged in genetic rescues compared to *Bckdk*^-/-^ mice (Figure 6M-O). Together, partial rescue of BCAA catabolic intermediates and resolution of initial intermediates of the TCA cycle are therefore achievable with *Dbt* haploinsufficiency in BCKDK deficiency.

*Dbt* haploinsufficiency also corrected elevations of key amino acids in the CNS, including lysine and glycine (Supplemental Figure 6C, F). Several additional amino acid markers of disease were similar in genetic rescue mice as compared to WT mice (Supplemental Figure 6G-I, K-M, O-R). Together these data support that there is partial restoration of amino acid levels, acylcarnitine levels, and organic acid levels through DBT haploinsufficiency in *Bckdk*^-/-^ mice.

## 4. Discussion

Pathogenic variants in BCKDK lead to a syndrome characterized by intellectual disability, postnatal microcephaly, autism, and seizures in human patients^4^. Previous models of *Bckdk* deficiency focused solely on the presence of seizures and hindlimb clasping as a phenotypic readout of *Bckdk* deficiency^8,17^. Therefore, in our study we characterized additional early onset, clinically-relevant neurological phenotypes and found significant neurodevelopmental delay and postnatal microcephaly in a recently published *Bckdk*^-/-^ mouse model^10^. From these assessments we can conclude that this mouse is a useful tool to investigate disease pathophysiology and assess potential therapies.

Since the discovery of BCKDK pathogenic variants, the mechanism of disease pathology was assumed to be hypercatabolism of BCAAs leading to their severe depletion. Our study revealed that there are multiple imbalances in BCAA catabolism and the TCA cycle due to BCKDK deficiency. First, we showed that there is accumulation of acylcarnitine derivatives downstream of the BCKDH step and decreased C5-OH levels further downstream suggesting a partial blockade in the BCAA catabolic pathway. Additionally, we found loss of BCKDK leads to imbalances in precursors to and intermediates of the TCA cycle with elevated pyruvate, aKG, fumarate, and malate levels but decreased succinate levels. Together, these data suggest that abnormal upstream BCAA metabolites induce imbalances in TCA cycle intermediates. These findings led us to question whether BCAA supplementation could both correct BCAA levels and normalize downstream TCA cycle changes.

Clinical studies of BCAA supplementation in patients with pathogenic variants of BCKDK are limited, and no randomized-control trials have been conducted. Available data suggest that in small cohorts early BCAA supplementation can partially improve disease phenotypes^5,7^.

Therefore, we systematically tested the efficacy of enteral BCAA supplementation in our mouse model of BCKDK deficiency. Our approach to replete BCAA levels was two-fold. The first was to test if peripheral administration would be sufficient to replete BCAA levels in the brain, since this had not yet been systematically proven^5,7^. Secondly, we aimed to increase the frequency of BCAA dosage to maximize repletion of BCAAs. Clinical studies revealed that multiple injections with BCAA supplementation (up to 6x) a day would be required, and even then, levels were insufficiently maintained within physiological ranges for most of the day^7^. Therefore, BCAA supplementation through dams via lactation was the best option to maximize frequent dosing of pups during the developmental timeframe without negatively impacting simultaneous neurodevelopmental assessments. In order to supplement BCAAs without them precipitating out of solution or reducing pup intake, we used a published protocol utilizing a 2% supersaturated solution of a ratio of BCAAs (2 Leu:1 Iso:1 Val) to maximize supplementation to pups^15^. BCAA supplementation did increase brain weight and size in *Bckdk*^-/-^ mice, likely through BCAAs known role to increase protein synthesis required for tissue growth^8^. However, BCAA supplementation did not rescue neurodevelopmental deficits, prevent postnatal microcephaly, or drastically correct biochemical abnormalities despite increasing systemic BCAA levels. The only correction BCAAs made to the BCAA catabolic pathway in *Bckdk*^-/-^ mice was to reduce pooling of the acylcarnitine derivatives one step below BCKDH.

These results reveal limitations in the ability of BCAA supplementation to rescue neurodevelopmental and biochemical changes in BCKDK deficiency. First, enteral administration of BCAAs does not significantly increase BCAA levels in the cerebral spinal fluid of patients^7^ nor in the brain of *Bckdk*^-/-^ mice. Therefore, restoration of BCAAs in the brain is not likely achievable through a peripheral administration route. Second, enteral supplementation may be insufficient to sustainably increase BCAAs in the periphery, as seen in patients where levels dropped to below the reference range 3-5 hours after dosing and in mice where leucine levels trended toward an increase but demonstrated great variability (Figure 3B)^7^.

As BCAA supplementation was insufficient to normalize disease biomarkers and neurodevelopment, we explored alternative therapeutic approaches. Under normal physiological conditions BCKDK serves to decrease activity of the BCKDH complex^8^. Therefore, we sought to mimic its inhibitory function by reducing expression of DBT, a key component of the BCKDH complex. Specifically, we tested if DBT haploinsufficiency, could normalize BCAA pathway and TCA cycle intermediates in the absence of BCKDK. Interestingly, early genetic re-regulation of BCAA catabolism, through *Dbt* haploinsufficiency, partially rescued microcephaly, behavioral phenotypes and restored most biochemical changes in *Bckdk*^-/-^ mice. Although intervention timing was not directly investigated in this work, our genetic rescue study supports that earlier intervention can affect outcomes such as survival and developmental phenotypes, both of which were improved with *Dbt* haploinsufficiency in *Bckdk*^-/-^ mice. In genetic rescue mice, *Dbt* mRNA transcript levels were only reduced by 32%, suggesting that future directions of this work could aim to further reduce these levels to potentially maximize phenotypic outcomes.

Still, caution should be taken to not fully deplete DBT levels in BCKDK deficiency, as the loss of DBT function would lead to a Maple-Syrup Urine Disease (MSUD), which induces severe neurological dysfunction^2^. *Dbt* haploinsufficiency may modulate this pathway by additional mechanisms by reducing BCAT activity, as they have been shown to interact in a metabalon^24^. This further illustrates that BCAA catabolism is fine-tuned to maintain metabolic homeostasis and neurodevelopment. Interestingly, not all changes in the downstream TCA cycle were changed by *Dbt* haploinsufficiency in *Bckdk*^-/-^ mice, suggesting the changes downstream of succinyl-CoA within the TCA cycle are inherent to absence of *Bckdk*. Previous studies have shown that pharmacologic inhibition of BCKDK with BT2 affected additional protein targets beyond BCKDH^25^. Therefore, a key future direction of this research is to investigate the impact of BCKDK in regulating the TCA cycle and overall macronutrient metabolism beyond BCAA catabolism.

## 5. Conclusion

In summary, we found that regulation of the rate of BCAA catabolism is essential to maintain metabolic homeostasis and neurodevelopment. As universal newborn screening has been transformative in other BCAA disorders, as is the case for MSUD^26^, the possibility of screening for BCKDK deficiency has been raised^7^. For screening to be effective, we must have therapeutics that improve clinical phenotypes and treat the underlying disease pathophysiology. Early studies of BCAA supplementation in patients have yielded some signs of benefit, yet outcomes vary and neurologic deficits remain in most if not all patients^7^. Here we also found that early BCAA supplementation does not fully correct and even worsens some disease phenotypes in a *Bckdk^-/-^* mouse model. In addition, our data suggest that decreasing flux through the BCAA pathway may provide an alternative therapeutic approach. Future, additional, well-controlled studies of BCAA supplementation and additional potential therapies are needed to improve outcomes in BCKDK deficiency.

## Acknowledgements

The *Bckdk*^-/-^ mouse was a gift from the Zoltan Arany lab at the University of Pennsylvania. The PCMD MicroCT Imaging Core is supported by Penn Center for Musculoskeletal Disorders (NIH/NIAMS P30 AR069619). We thank the Penn Metabolomics Core (RRID:SCR_022381) in the Cardiovascular Institute at the University of Pennsylvania for metabolomics analyses. The data for this manuscript was generated in the Penn Cytomics and Cell Sorting Shared Resource Laboratory at the University of Pennsylvania and is partially supported by the Abramson Cancer Center NCI Grant (P30 016520). The research identifier number is RRid:SCR_022376. We would like to thank Adele Harman, Technical Director of the Children’s Hospital of Philadelphia Transgenic Core Facility for creating the *Dbt* mouse line.

## 7. Author contributions

R.A. and Z.A. provided scientific direction. L.O. and R.A. designed the study. L.O., A.K., R.P., M.N., and E.D. carried out experiments. L.O. collected samples, performed biochemical experiments, data analysis, and interpretation. M.N. executed a subset of biochemical experiments and data analyses, and L.O. collaboratively interpreted the data. L.O. coordinated neurodevelopmental studies, A.K. and R.P. executed them. L.O. and A.K. statistically analyzed them with R.A.’s advisal. N.P. assisted with data entry and quality assurance of genetic rescue developmental studies. E.D. performed µCT studies. E.D. and L.O. analyzed the µCT data collaboratively. L.O. wrote the manuscript with input from all authors. R.A. and Z.A. initiated and L.O. facilitated collaborations. R.A. and Z.A. allocated funding for this study.

## 8. Data Availability

Data generated and analyzed from this study are included in this article and the supplementary information. Data supporting these studies findings are available from the corresponding author with reasonable request.

