## Supplemental Figures and Tables for "Partial suppression of BCAA catabolism as a potential therapy for BCKDK deficiency"

### 1. Supplementary Figures

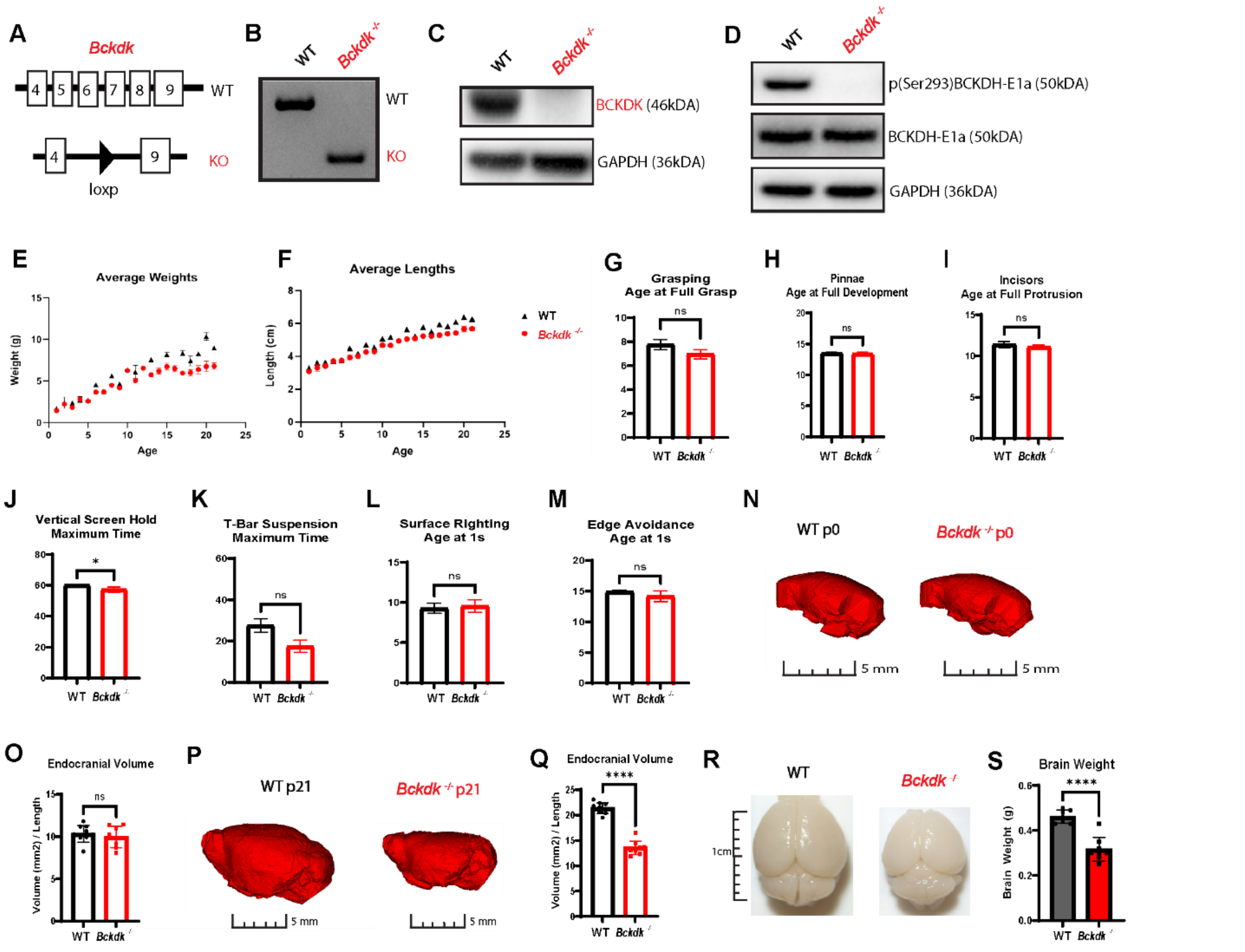


***Supplementary Figure 1 Biochemical validation, insignificant neurodevelopmental changes, and reduced brain size.*** *A sex balanced cohort of WT (n=27) and Bckdk$^{-/-}$ (n=21) mice were compared for neurodevelopmental assessment. An unpaired t-test was used for statistical comparison. In all panels, data are represented as mean with s.e.m. (* $p < 0.05$, ** $p<0.005$, *** $p < 0.001$, **** $p<0.0001$, ns=non-significant). (A) Bckdk WT and KO allele graphically illustrating loss of exon 5-8 in the KO allele. (B) Genotyping Results of Bckdk PCR discerning WT from KO mice. (C) Absence of BCKDK in Bckdk$^{-/-}$ mice as seen by western blot. (D) Absence of pBCKDH-E1a but not BCKDH-E1a in Bckdk$^{-/-}$ mice as seen by western blot. (E) Reduced average body weight in Bckdk$^{-/-}$ mice diverging from WT mice at 2 weeks of age and reducing in the 3$^{rd}$ week of age. (F) Insignificantly altered body length in Bckdk$^{-/-}$ mice relative to WT mice. (G) Grasping was not significantly altered in Bckdk$^{-/-}$ mice compared to WT mice. (H) Lack of changes in pinnae development in Bckdk$^{-/-}$ mice compared to WT mice. (I) No difference in the age of full protrusion of incisors in Bckdk$^{-/-}$ mice compared to WT mice. (J) Reduced maximum hold time on the vertical screen hold in Bckdk$^{-/-}$ mice compared to WT mice. (K) Trend decrease in maximum time on T-bar suspension in Bckdk$^{-/-}$ mice. (L) No changes in age of surface righting. (M) No difference in the age of edge avoidance. (N) Reduced brain volume in Bckdk$^{-/-}$ mice compared to WT mice as seen by representative image of 3D reconstruction from µCT scans. (O) Significantly decreased endocranial volume in Bckdk$^{-/-}$ mice. (P) Reduced brain size in Bckdk$^{-/-}$ mice compared to WT mice visualized by representative images of mouse brains. (Q) Drastically reduced brain weight in Bckdk$^{-/-}$ (n=8) mice compared to WT (n=7) mice.*

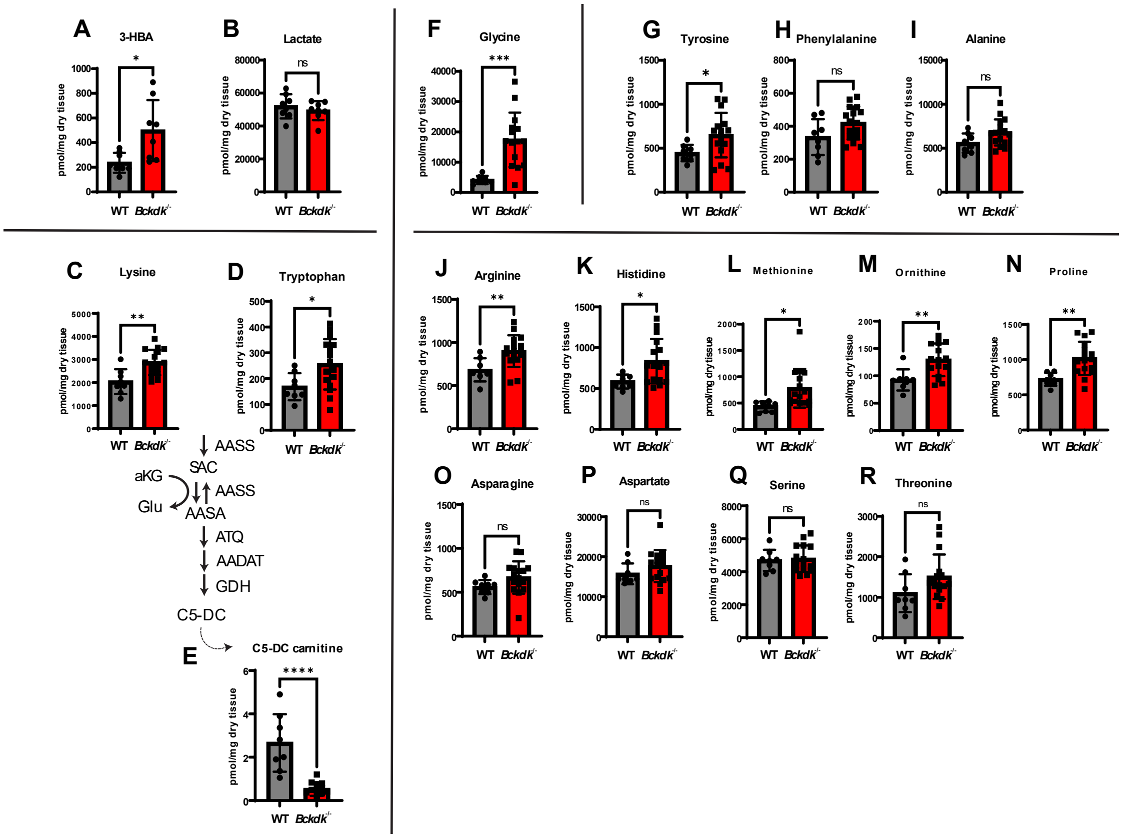

***Supplementary Figure 2 Amino acids and additional acylcarnitine levels in the brain of WT and Bckdk$^{-/-}$ mice at p21.*** *A sex balanced cohort of WT (n=8) and Bckdk$^{-/-}$ (n=15) mice were sent for amino acid and acylcarnitine analysis, and a subset of Bckdk$^{-/-}$ mice (n=7) for organic acids measured by mass spectrometry. Unpaired t-tests were performed to compare the two genotypes. In all panels, data are represented as mean with s.e.m. (* $p < 0.05$, ** $p<0.005$, *** $p < 0.001$, **** $p<0.0001$, ns=non-significant). (A) Elevated 3-HBA levels in Bckdk$^{-/-}$ mice, the metabolite downstream of leucine degradation. (B) Insignificant changes in lactate levels. (C) Accumulation of lysine in Bckdk$^{-/-}$ mice. (D) Increased levels of tryptophan in Bckdk$^{-/-}$ mice. (E) Depletion of downstream lysine metabolite glutaryl-carnitine (C5-DC carnitine) in Bckdk$^{-/-}$ mice. (F) Significantly elevated glycine levels in Bckdk$^{-/-}$ mice relative to WT levels. (G) Elevation of tyrosine in Bckdk$^{-/-}$ mice, a clinical marker of BCKDK deficiency. (H) Trend increase in phenylalanine in Bckdk$^{-/-}$ mice, another clinical marker. (I) Trend elevation of alanine in Bckdk$^{-/-}$ mice, another clinical marker. (J) Significant increase in arginine levels in Bckdk$^{-/-}$ mice. (K) Elevation of histidine levels in Bckdk$^{-/-}$ mice. (L) Higher levels of methionine in Bckdk$^{-/-}$ mice. (M) Increased levels of ornithine in Bckdk$^{-/-}$ mice. (N) Higher proline levels in Bckdk$^{-/-}$ mice. (O) Trend increase in asparagine levels in Bckdk$^{-/-}$ mice. (P) Trend elevation of aspartate levels in Bckdk$^{-/-}$ mice. (Q) Insignificant changes in serine levels in Bckdk$^{-/-}$ mice. (R) Trend accumulation of threonine in Bckdk$^{-/-}$ mice.*

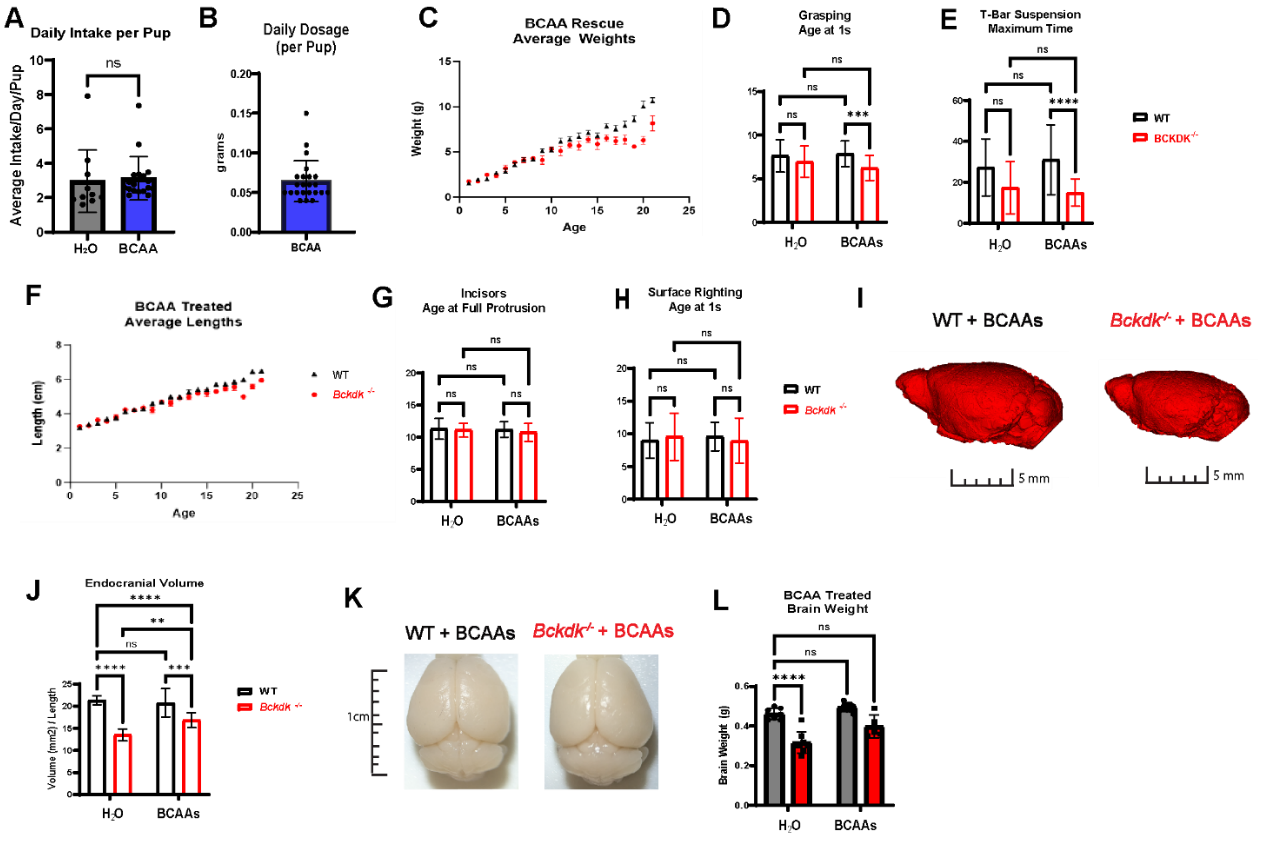


***Supplementary Figure 3 BCAA supplementation further exacerbates neurodevelopmental differences but increases brain size in Bckdk$^{-/-}$ Mice.*** *A sex balanced cohort of WT (n=39) and Bckdk$^{-/-}$ (n=28) mice were compared for all neurodevelopmental assessments. 2-way ANOVA with multiple comparisons was used for all statistical analyses. In all panels, data are represented as mean with s.e.m. (* $p < 0.05$, ** $p<0.005$, *** $p < 0.001$, **** $p<0.0001$, ns=non-significant). (A) Similar daily intake per pup revealing self-regulation of BCAA dosage with litter size in lactating dams ($H_2O$(n=20), BCAA (n=24)). (B) Average dosage per day per pup in BCAA administrated lactating dams (n=24). (C) Reduced body weight at 2 weeks in Bckdk$^{-/-}$ mice compared to WT mice regardless of BCAA intervention. (D) BCAAs further exacerbated differences in grasping in Bckdk$^{-/-}$ and WT mice. (E) Worsening of maximum time on T-bar in BCAA treated Bckdk$^{-/-}$ and WT mice. (F) Insignificant alterations in average length of BCAA treated Bckdk$^{-/-}$ mice. (G) Incisor eruption is not significantly impacted by genotype or treatment group. (H) Surface righting was not significantly impacted by genotype nor BCAA treatment. (I) Reduced brain volume in BCAA treated Bckdk$^{-/-}$ mice compared to BCAA treated WT mice as seen by representative image of 3D reconstruction from µCT scans. (J) Significantly decreased endocranial volume in BCAA treated Bckdk$^{-/-}$ mice. (K) Reduced brain size in BCAA treated Bckdk$^{-/-}$ mice compared to BCAA treated WT mice visualized by representative images of mouse brains. (L) Partial rescue of brain weight in BCAA treated Bckdk$^{-/-}$ (n=5) mice with insignificant changes from untreated WT (n=7) brain weights. Also compared are BCAA treated WT mice (n=9), and untreated Bckdk$^{-/-}$ mice (n=8).*

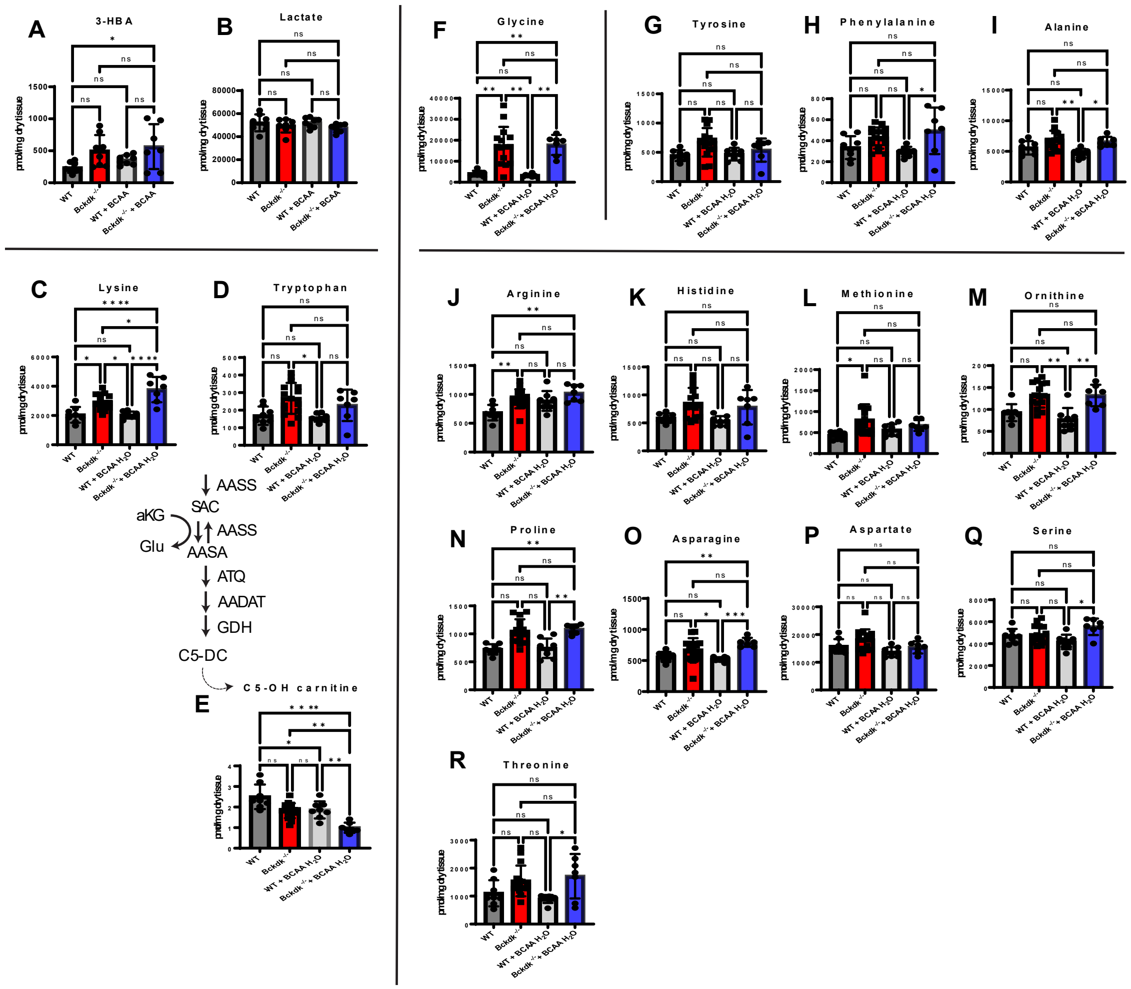


***Supplementary Figure 4 Insignificant alterations of amino acids and acylcarnitine changes in BCAA treated Bckdk$^{-/-}$ mice at p21.*** *A sex balanced cohort of BCAA treated WT (n=8) and Bckdk$^{-/-}$ (n=7) mice were compared to untreated WT (n=8) and Bckdk$^{-/-}$ (n=15) data from figure 2. Brain amino acid, acylcarnitine, and organic acid levels were measured by mass spectrometry. 2-way ANOVAs were performed to compare all four groups for genotype and BCAA intervention differences. In all panels, data are represented as mean with s.e.m. (* $p < 0.05$, ** $p<0.005$, *** $p < 0.001$, **** $p<0.0001$, ns=non-significant). (A) Further elevation of 3-HBA levels in BCAA treated Bckdk$^{-/-}$ mice. (B) Insignificant changes in lactate levels regardless of treatment or genotype. (C) Further accumulation of lysine in BCAA treated Bckdk$^{-/-}$ mice compared to Bckdk$^{-/-}$ mice, both relative to WT levels. (D) Similar tryptophan levels in BCAA treated Bckdk$^{-/-}$ mice relative to WT levels. (E) Significant depletion of downstream lysine metabolite glutaryl-carnitine (C5-DC carnitine) in BCAA treated Bckdk$^{-/-}$ mice. (F) Significantly elevated glycine levels in BCAA treated Bckdk$^{-/-}$ mice relative to untreated and treated WT levels. (G) Insignificant changes in tyrosine in BCAA treated Bckdk$^{-/-}$ mice, a clinical marker of BCKDK deficiency. (H) Increase in phenylalanine levels in BCAA treated Bckdk$^{-/-}$ mice compared to BCAA treated WT mice. (I) Elevation of alanine levels in BCAA treated Bckdk$^{-/-}$ mice, compared to BCAA treated WT mice. (J) Similar increase in arginine levels in BCAA treated Bckdk$^{-/-}$ mice compared to untreated BCKDK$^{-/-}$ mice. (K) Trend elevation of histidine levels in BCAA treated Bckdk$^{-/-}$ mice compared to untreated WT mice. (L) Lower levels of methionine in BCAA treated Bckdk$^{-/-}$ mice compared to untreated BCKDK$^{-/-}$ mice. (M) Further exacerbation of increased ornithine levels in BCAA treated Bckdk$^{-/-}$ mice relative to BCAA treated WT mice. (N) Significantly higher proline levels in BCAA treated Bckdk$^{-/-}$ mice compared to untreated WT mice. (O) Higher increase in asparagine levels in BCAA treated Bckdk$^{-/-}$ mice relative to untreated WT mice. (P) Minimal changes in aspartate levels in BCAA treated Bckdk$^{-/-}$ mice compared to all groups. (Q) Further exacerbation of elevated serine levels in BCAA treated Bckdk$^{-/-}$ mice relative to BCAA treated WT mice. (R) Further accumulation of threonine in BCAA treated Bckdk$^{-/-}$ mice compared to BCAA treated WT mice.*

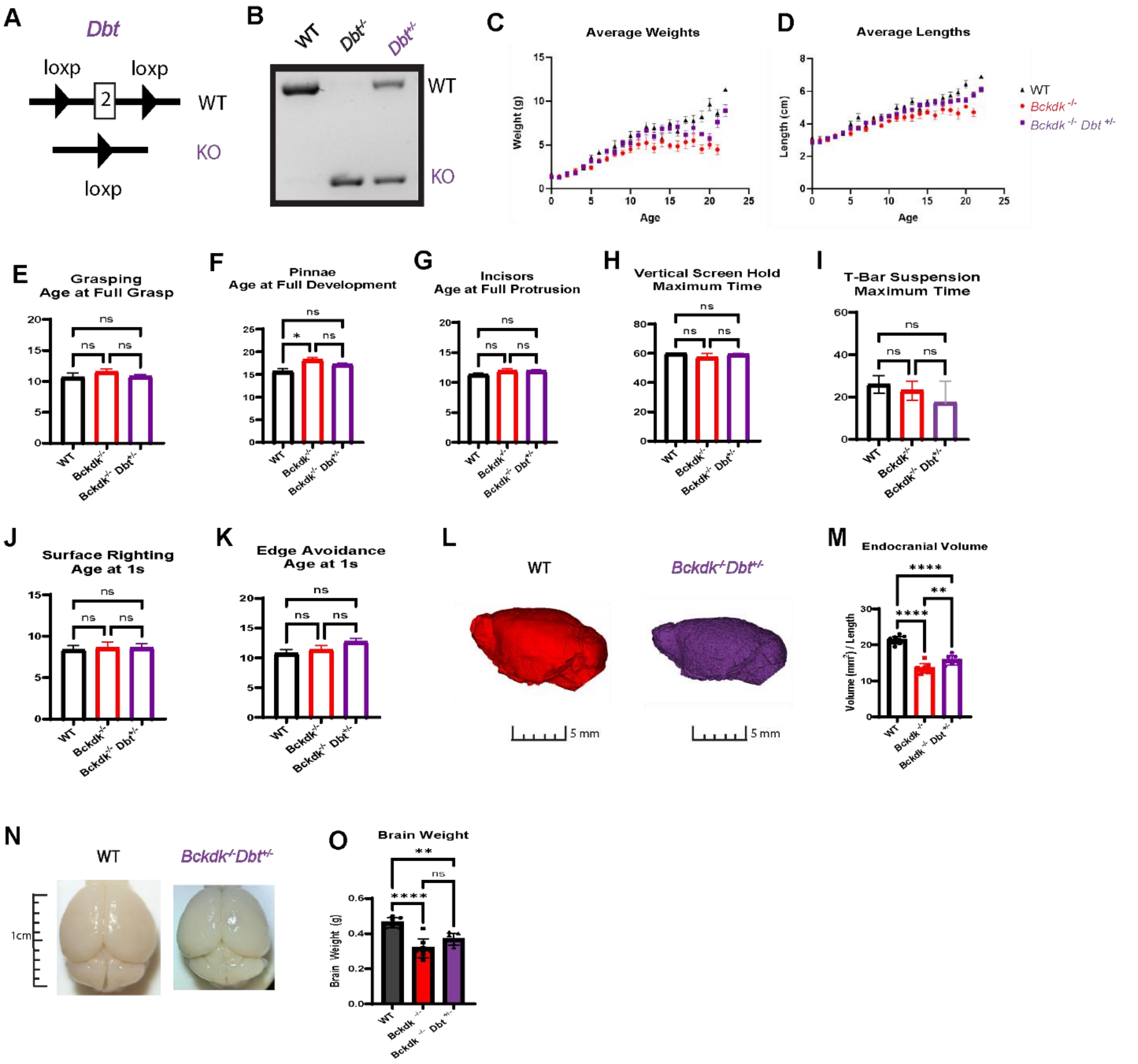


***Supplementary Figure 5 Genetic rescue mice have partial rescue of neurodevelopmental delay and microcephaly.*** *A sex balanced cohort of WT (n=12), Bckdk$^{-/-}$ (n=13), and Bckdk$^{-/-}$Dbt$^{+/-}$ (n=26) mice were compared for all neurodevelopmental assessments. One-way ANOVAs with multiple comparisons were used for comparisons between the three groups. In all panels, data are represented as mean with s.e.m. (* $p < 0.05$, ** $p<0.005$, *** $p < 0.001$, **** $p<0.0001$, ns=non-significant). (A) Expected product size of PCR products for genotyping of Dbt WT band (621 bp) and KO band (311 bp). (B) Genotyping Results of DBT PCR discerning WT from KO alleles. (C) Partial rescue of body weight gain over time in Bckdk$^{-/-}$Dbt$^{+/-}$ mice. (D) Average length of mice was not significantly altered in all groups over time. (E) Grasping is similar in Bckdk$^{-/-}$Dbt$^{+/-}$ mice compared to WT and Bckdk$^{-/-}$ mice. (F) Rescue of age at full pinnae development in Bckdk$^{-/-}$Dbt$^{+/-}$ relative to WT mice. (G) Insignificant changes in age of incisor eruption regardless of genotype. (H) No drastic changes in maximum time on vertical screen hold in all groups. (I) Trend decrease in maximum time on T-bar suspension in Bckdk$^{-/-}$Dbt$^{+/-}$ compared to WT and Bckdk$^{-/-}$ mice. (J) Insignificant changes in age of surface righting in Bckdk$^{-/-}$Dbt$^{+/-}$ compared to WT and Bckdk$^{-/-}$ mice. (K) No significant changes in edge avoidance in Bckdk$^{-/-}$Dbt$^{+/-}$ compared to WT and Bckdk$^{-/-}$ mice. (L) Partially rescued brain volume in Bckdk$^{-/-}$Dbt$^{+/-}$ compared to WT mice as seen by representative image of 3D reconstruction from µCT scans. (M) Partially restored endocranial volume in Bckdk$^{-/-}$Dbt$^{+/-}$ mice compared to Bckdk$^{-/-}$ mice, both relative to WT mice. (N) Similar brain size in Bckdk$^{-/-}$Dbt$^{+/-}$ mice compared to WT mice visualized by representative images of mouse brains. (O) Incrementally increased average brain weight of Bckdk$^{-/-}$Dbt$^{+/-}$ (n=7) mice compared to WT (n=7) mice. Also compared is Bckdk$^{-/-}$ mice (n=8).*

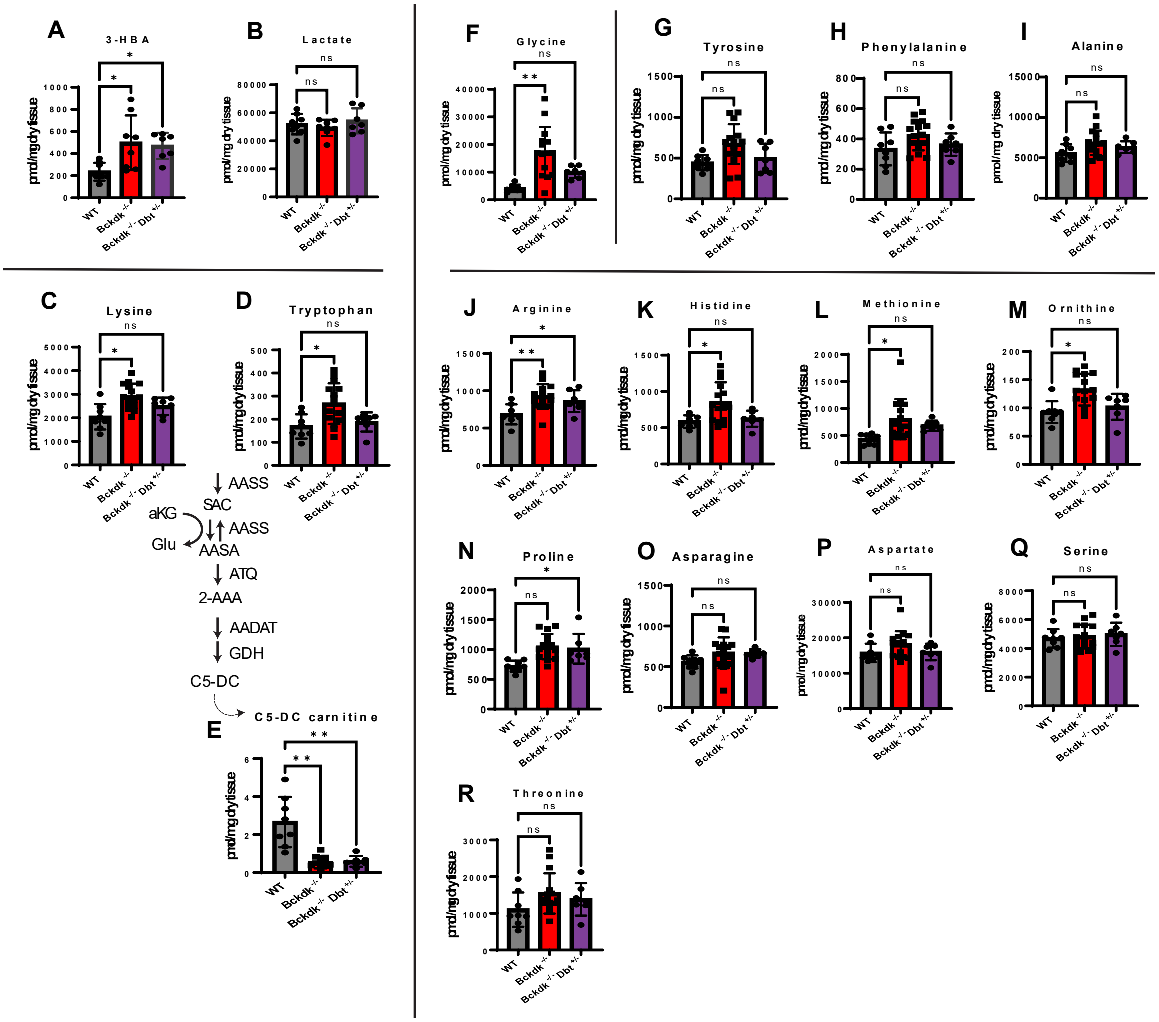


***Supplementary Figure 6 Rescue of amino acids and acylcarnitine levels in the brain Bckdk$^{-/-}$Dbt$^{+/-}$ mice.*** *A sex balanced cohort of Bckdk$^{-/-}$Dbt$^{+/-}$ (n=7) mice were compared to untreated WT (n=8) and Bckdk$^{-/-}$ (n=15) metabolic data from the original characterization shown in Figure 2. Amino acid, acylcarnitine, and organic acids levels were measured by mass spectrometry. One-way ANOVA with multiple comparisons was used to compare genotypes. In all panels, data are represented as mean with s.e.m. (* $p < 0.05$, ** $p<0.005$, *** $p < 0.001$, **** $p<0.0001$, ns=non-significant). (A) Similar elevation of 3-HBA levels in Bckdk$^{-/-}$Dbt$^{+/-}$ mice compared to WT levels. (B) Insignificant changes in lactate levels regardless of genotype. (C) Resolution of lysine levels in Bckdk$^{-/-}$Dbt$^{+/-}$ mice similar to WT levels. (D) Similar tryptophan levels in Bckdk$^{-/-}$Dbt$^{+/-}$ mice relative to WT mice. (E) Significant depletion of downstream lysine metabolite glutaryl-carnitine (C5-DC carnitine) in Bckdk$^{-/-}$Dbt$^{+/-}$ mice compared to WT mice. (F) Rescue of glycine levels in Bckdk$^{-/-}$Dbt$^{+/-}$ mice back to nearly WT levels. (G) Insignificant changes in tyrosine levels in Bckdk$^{-/-}$Dbt$^{+/-}$ mice compared to WT levels, which is a clinical marker of BCKDK deficiency. (H) Similar phenylalanine levels Bckdk$^{-/-}$Dbt$^{+/-}$ mice compared to WT mice. (I) Same alanine levels in Bckdk$^{-/-}$Dbt$^{+/-}$ mice relative to WT mice. (J) Partial rescue of arginine levels in Bckdk$^{-/-}$Dbt$^{+/-}$ mice compared to Bckdk$^{-/-}$ and WT mice. (K) Resolution of histidine levels in Bckdk$^{-/-}$Dbt$^{+/-}$ back to WT levels. (L) Insignificant changes in methionine levels in Bckdk$^{-/-}$Dbt$^{+/-}$ mice relative to WT mice. (M) Rescue of ornithine levels in Bckdk$^{-/-}$Dbt$^{+/-}$ back to WT levels. (N) Elevated proline levels in Bckdk$^{-/-}$Dbt$^{+/-}$ mice compared to WT mice. (O) Insignificant changes in asparagine levels in Bckdk$^{-/-}$Dbt$^{+/-}$ mice compared to WT levels. (P) Minimal changes in aspartate levels in Bckdk$^{-/-}$Dbt$^{+/-}$ compared to all groups. (Q) No changes in serine levels in Bckdk$^{-/-}$Dbt$^{+/-}$ mice relative to all groups. (R) Trend increase in threonine levels in Bckdk$^{-/-}$Dbt$^{+/-}$ mice compared to WT levels.*

#### 13. Supplementary Tables

Supplementary Table 1. The list of primers and sequences for real-time PCR.

| Gene | Forward Primer | Reverse Primer |
|---|---|---|
| *Dbt* | CACGTCACTCCCTGAGAACA | CGAATTCCTTCTCCGATGTC |
| *Actin* | CTGTATTCCCCTCCATCGTG | CTCGTCACCCACATAGGAGTC |

Supplementary Table 2. The list of antibodies and dilutions used for Western Blot

| Antibody | Dilution | Catalog Number |
|---|---|---|
| BCKDK | 1:1000 | Ab128935 |
| pBCKDH-E1a (Ser293) | 1:1000 | CST 40368 |
| BCKDH-E1a | 1:2000 | CST 90198 |
| GAPDH | 1:5000 | Proteintech 60004-1-Ig |
